# IGLoo: Profiling the Immunoglobulin Heavy chain locus in Lymphoblastoid Cell Lines with PacBio High-Fidelity Sequencing reads

**DOI:** 10.1101/2024.07.20.604421

**Authors:** Mao-Jan Lin, Ben Langmead, Yana Safonova

## Abstract

New high-quality human genome assemblies derived from lymphoblastoid cell lines (LCLs) provide reference genomes and pangenomes for genomics studies. However, the characteristics of LCLs pose technical challenges to profiling immunoglobulin (IG) genes. IG loci in LCLs contain a mixture of germline and somatically recombined haplotypes, making them difficult to genotype or assemble accurately. To address these challenges, we introduce `IGLoo`, a software tool that implements novel methods for analyzing sequence data and genome assemblies derived from LCLs. `IGLoo` characterizes somatic V(D)J recombination events in the sequence data and identifies the breakpoints and missing IG genes in the LCL-based assemblies. Furthermore, `IGLoo` implements a novel reassembly framework to improve germline assembly quality by integrating information about somatic events and population structural variantions in the IG loci. We applied `IGLoo` to study the assemblies from the Human Pangenome Reference Consortium, providing new insights into the mechanisms, gene usage, and patterns of V(D)J recombination, causes of assembly fragmentation in the IG heavy chain (IGH) locus, and improved representation of the IGH assemblies.

## 1 Introduction

Immunoglobulin (IG) gene loci are essential for the development of B-cell receptors (BCRs) and antibodies (Abs), which are one of two main components of the adaptive immune system. Most mammalian genomes contain three IG loci: one immunoglobulin heavy chain (IGH) locus and two immunoglobulin light chain loci, kappa chain (IGK) and lambda chain (IGL). The IGH locus comprises a series of variable (V), diversity (D), joining (J), and constant (C) genes, while the IGK and IGL loci have a similar structure but without the D genes. During B-cell maturation, a process known as V(D)J recombination occurs, where one V, D (in case of IGH locus), and J gene are randomly selected and joined to form a rearranged V(D)J segment. In the IGH, D-J recombination precedes V-D recombination, resulting in the excision of the sequence between the selected D and J genes, followed by the excision of the sequence between the selected V gene and the newly formed D-J complex (1) (2). These rearrangements, processed through double-strand breaks and subsequent repair at the recombination signal sequences (RSSs) flanking the selected IG genes, generate a diverse antibody repertoire and mount the potential to bind and neutralize a wide range of antigens. The genotype of these loci shape an individual’s antibody response, highlighting the importance of the germline diversity of IG genes and loci in adaptive immunity (3) (4) (5).

Lymphoblastoid cell lines (LCLs) are a widely used system for providing a continuous source of human cells. LCLs are easy to prepare and maintain, and have a low somatic mutation rate in continuous culture (6). LCLs can provide an ongoing source of DNA that is almost identical to original normal cells when compared using whole genome sequencing (WGS) (7). LCLs were used in several major consortium projects that surveyed human genetic variation, such as the 1000 Genomes Project (8), HapMap (9), Genome in a Bottle (10), and the Human Pangenome Reference Consortium (HPRC) (11).

On the other hand, previous studies also showed that the DNA derived from LCLs contains V(D)J recombinations and somatic hypermutation (SHMs) events (12) affecting IG loci and making them harder to genotype and assemble (13). This is because LCLs are generated by transforming B-cells using Epstein-Barr Virus (EBV). The original B-cells likely already harbored V(D)J recombination events, as normal B-cells typically do.

There have been a few attempts to profile human IG loci. Rodriguez et al (14) and Gibson et al (15) used PacBio single molecule high-fidelity (HiFi) sequencing to target and study IG loci from LCLs. The former focused on the IGH locus, while the latter examined the IGL locus. Both papers report the germline genotypes of the IG loci and annotate which regions on the loci are unknown due to V(D)J recombination. A more recent study by Rodriguez et al (3) circumvented these challenges by using sequence data from peripheral blood mononuclear cells (PBMCs) or polymorphonuclear leukocytes instead of LCLs thus avoiding the obstacles posed by V(D)J recombination. Nonetheless, LCLs are still widely used and easily accessible in existing database. As a result, there remains a significant need for an accurate tool for IG profiling in LCL data.

HPRC aims to build a human pangenome reference to represent the global genetic diversity. In their year 1 release, HPRC provided 47 phased, diploid assemblies from 15 subpopulations. All 47 samples are derived from LCLs. The samples were sequenced with PacBio HiFi sequencing with an average read depth of 39.7*×*, except HG002 which had average coverage *>* 130*×*. The WGS Illumina short reads of the individuals’ parents are also available to phase the individual’s assembly. The HiFi reads of the individual and Illumina reads of the parents were assembled into phased assemblies with the trio-binning mode of hifiasm (16). Though the assemblies are carefully curated, no particular measures have been employed to study the assembly quality in the IG loci and how they may have been affected by the LCL input. At this time, little is known about how representative the assemblies are of the germline IG sequence.

We present `IGLoo`, a toolkit for assessment and improvement of the IGH locus representation in LCL sequencing data. `IGLoo` profiles the somatic V(D)J recombination events in an LCL genome and measures their clonality. Then it improves the germline assembly of the IGH locus by removing the reads representing somatic haplotypes driven by V(D)J recombination and reassembling the dataset using hifiasm and MaSuRCA (17). Finally, `IGLoo` assesses the quality of IGH locus assembly by analyzing the breakpoints and missing IGH genes.

We applied `IGLoo` to WGS LCL datasets collected from 47 individuals in the HPRC year-1 release. We observed diverse levels of clonality, as well as varying quality of the assemblies in the IGH locus. By analyzing the HiFi reads, we also found evidence of a range of known and novel non-canonical V(D)J recombination events. To our knowledge, `IGLoo` is the first work to systematically analyze and report non-canonical recombination events in long-read datasets and assemblies. We further applied the `IGLoo` reassembly framework to improve the assembly result of IGH locus. On average, `IGLoo` reassembled IGH locus covered 10.8 more IGH genes per individual than HPRC year-1 assemblies.

## 2 Results

The `IGLoo` tool, which analyzes the IGH locus of human lymphoblastoid cell lines (LCLs) sequenced using PacBio HiFi reads, consists of three modules. The first module, `IGLoo --read`, identifies and quantifies V(D)J recombination events present in a sample’s read alignment, offering insights into the recombination event frequency and IGH gene usage. The second module, `IGLoo --asm`, evaluates an assembly by cataloging present or missing IGH genes and pinpointing regions where the assembly was fragmented due to breakpoints. Lastly, the `IGLoo --ReAsm` module utilizes the output from `IGLoo --read` to modify input reads and construct new *de novo* and reference-based assemblies, resulting in an improved assembly that more comprehensively captures the germline genome.

We applied the `IGLoo` modules to human LCL samples representing 47 individuals available at the HPRC (11). Each sample consists of WGS HiFi reads, phased genome assemblies, and parental illumina read data.

### 2.1 IGLoo --read profiles the V(D)J recombination events in LCLs

To identify the V(D)J recombination events in the WGS data, it is essential to consider the mechanism of the V(D)J recombination. V(D)J recombination initiates and terminates at RSSs flanking the selected V, D, and J genes. Therefore a read carrying a V(D)J recombination event represents a deletion from one specific RSS to another RSS. WGS HiFi sequencing data is particularly advantageous for this task because typical V(D)J recombinations span less than 500 bp, while non-canonical recombination events can span thousands of bps (Section 2.1.3). HiFi reads are long enough to cover these events while also providing sequence context on either side of the recombination. Moreover, WGS HiFi enables detection of unusually long recombination events, such as RSS skipping (18) and the LAIR-1 insertion (19) as well as discovery novel types of non-canonical recombination events.

To profile the V(D)J recombination events in a sample, the `IGLoo --read` module takes either read alignments (BAM/CRAM) or HiFi reads (FASTA) and collects the reads spanning recombination events. When a HiFi read overlaps a V(D)J recombination event, a read aligner like minimap2 (20) will generally yield an alignment with distinct segments of the read aligning to either side of the recombination event. We call this a split alignment, and we say the read is a split-aligned read. Since the IGH locus is highly polymorphic (14), and no standard human reference genome covered all IGH genes, `IGLoo`

`--read` utilizes a multi-reference strategy (Section 4.1) to maximize the number of HiFi reads showing evidence of V(D)J recombination. `IGLoo --read` then reports the best-supported recombination events based on alignments to three human reference genomes: GRCh37 (21), GRCh38 (22), and T2T-CHM13 (23). The profile of the recombination events is then used to compute the gene usage and clonality of the cell line.

`IGLoo --read` scans and analyzes only the reads fully or partially mapped to the region stretching from J gene to D gene. In this way, we can detect reads that are split-aligned due to V-D and D-J recombination, or a combination of both. This way, reads that are split-aligned because of structural variations (SV) in the V gene locus will likely not be included in the analysis.

Since V(D)J recombination initiates and terminates at the RSSs of the selected V/D/J genes, when a read is split-aligned to two IGH genes due to such an event, we anticipate the split sites to be in close proximity to the RSSs. Alongside recombination-driven deletions, non-genomic nucleotides contributing to the diversity at the IG gene junctions and somatic hypermutations (SHMs) can also take place on the recombined fragment. However, due to the length of HiFi reads, the impact of the junctional diversity and SHMs on the analysis is minimal. We assessed the distance between split sites and the nearest V, D, J RSS for all split-aligned reads. Of the 1, 730 split-aligned reads across the 47 samples, totaling 3, 357 split-segments with some alignments split into more than two segments, over 93% of segments were within 50 bp of an RSS (Figure S1). Therefore, we considered split-aligned pairs where the split position is within 50 bp of the RSS as “confident” events and focused our gene usage analysis solely on these ”confident” events.

#### 2.1.1 Gene usage across cell lines

To detect the IGH gene usage frequency, we analyzed the HiFi reads that showed evidence of V(D)J recombination events and collected the respective V, D, or J genes used in these events. In Figure 1 **a**, we show the J and V gene pair usage in “confident” canonical V(D)J recombination events and in **b** the J and D gene usage for “confident” D-J only recombination, i.e. partial recombination, events across the 47 samples. The non-canonical recombination events are also included in the Figure 1 **b**, where only the first D-J gene pair is counted. A recombination event is counted only once per individual. The most widely used J gene is *IGHJ4* in both complete V(D)J and partial D-J recombination events, the same as the result in (3). In D-J recombination events, the most widely used D gene was *IGHD3-22*, which is consistent with the analysis of expressed V(D)J recombinations (Rep-Seq) described in (18). For complete V(D)J recombination events, the most widely used genes were *IGHV3-23* and *IGHV3-33*, which is very similar to the gene usage analysis using longs read for both isotype IgM and IgG described by Rodriguez *et al* (3).

**Figure 1:**
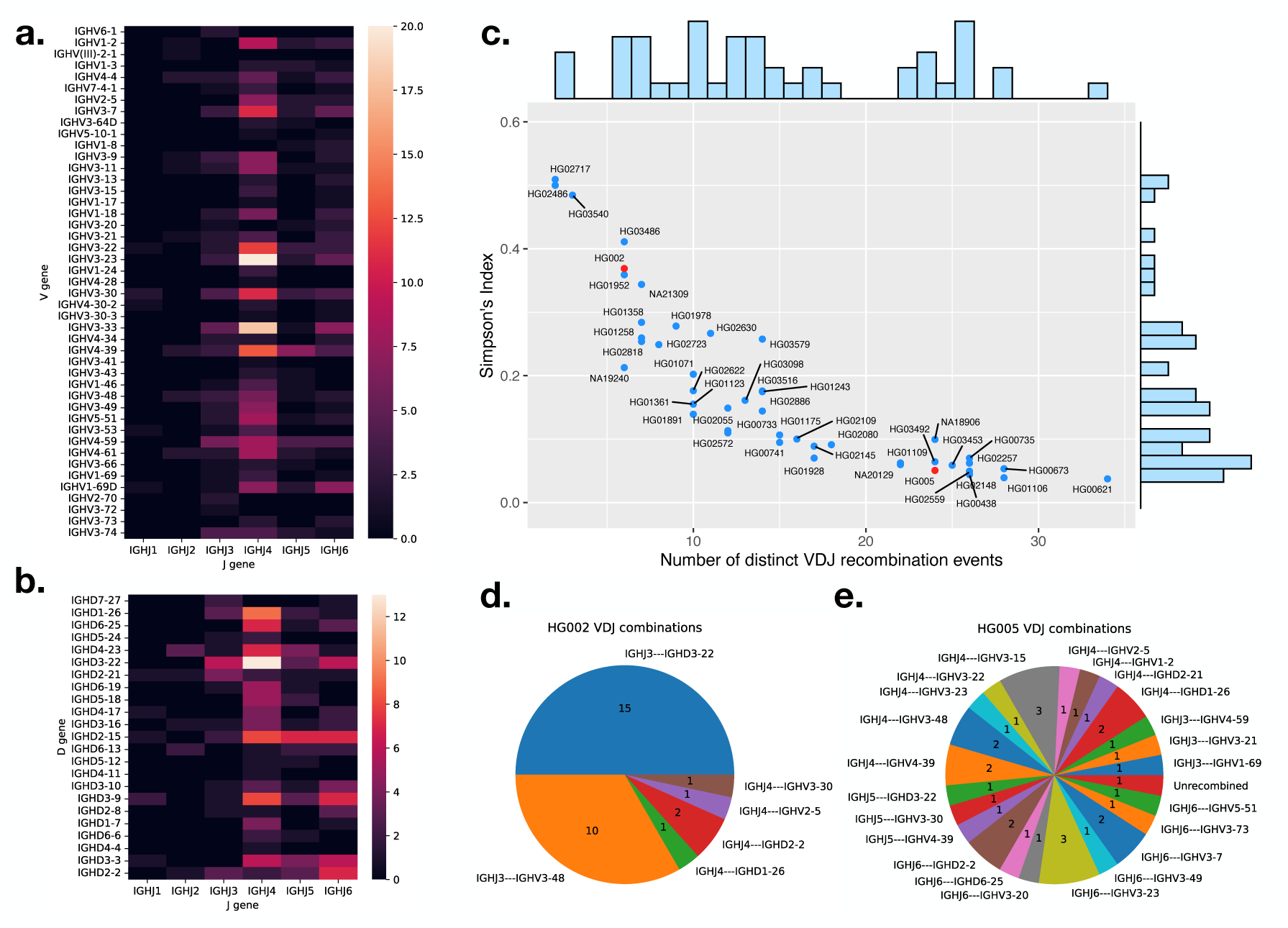
**a.** gene usage heatmap between V and J genes in complete V(D)J recombination. **b.** gene usage heatmap between J and D genes in partial recombinations and non-canonical recombinations. **c.** number of unique recombination event in a sample versus the Simpson’s index of the reads supporting each clone. The positions of HG002 and HG005 are highlighted in red. **d.** unique recombination events and number of reads supporting the event for cell line HG002 and **e.** cell line (HG005).

Our analysis also revealed gene usage patterns that are challenging to detect using Rep-Seq data. The first involves an 11 nt long D gene *IGHD7-27* that is difficult to distinguish from random matches in Rep-Seq data. In the LCL data, the D gene *IGHD7-27* was detected and found to participate in partial V(D)J recombination events with *IGHJ6* and *IGHJ3*. Additonally, we identified four pseudo V genes – *IGHV(III)-2-1*, *IGHV1-17*, *IGHV3-22*, and *IGHV3-41* – participating in canonical V(D)J recombination events. Contrary to the classification as pseudogenes in the IMGT database (24), *IGHV3-22* is used in 21 different recombination events across the samples, while recombination events involving *IGHV(III)-2-1*, *IGHV1-17*, and *IGHV3-41* were each detected in only one sample. In all cases, the split sites of the recombination events are both within 5 bases of the corresponding RSS (Figure S2, S3, S4). Although a previous study (14) did not show any usage of the four pseudogenes, we demonstrated that they can actually be functional V genes with low usages.

#### 2.1.2 Clonality of the cell lines

We collected all the V(D)J recombination events of each sample, and used Simpson’s index (SI) to measure the clonality of the sample. This is calculated as:

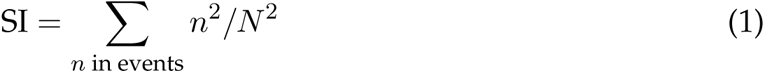

where *n* is the number of reads in a unique V(D)J recombination event, and *N* is the total number of reads containing V(D)J recombination event. The higher the SI, the more monoclonal the sample is.

We plot all the 47 samples according to their number of distinct V(D)J recombination events and SI in Figure 1 **c**. The negative trend is expected, since SI will tend to decrease with increasing number of distinct events. The Pearson’s correlation coefficient *r* between SI and the number of distinct events is *−*0.843 in this case. For a monoclonal sample, we expect SI to be near 0.5, representing one dominant recombination event per haplotype. This is the case for a few samples that appear toward the left of Figure 1 **c**. Most samples, though, have SI less than 0.5, indicating a degree of polyclonality. There was no clear separation between the SI values for monoclonal and polyclonal samples; rather, they formed a continuous spectrum. Note that our method will generally underestimate polyclonality due to the finite depth of WGS HiFi samples. For instance, the average coverage in these samples was about 38*×* in the IGH locus, impairing our ability to measure SI below about 0.03.

To illustrate the differences between monoclonal and polyclonal samples, we picked one sample (HG002) situated close to the monoclonal end and one sample (HG005) situated close to the polyclonal end for comparison. The positions of the two samples are highlighted in red in Figure 1 **c**, and their V(D)J recombination events are shown in Figure 1 **d** and **e**, respectively. The majority of the events for HG002 fall into two categories, consistent with this being a cell line with two haplotypes, each with a dominant recombination event. For HG005, on the other hand, the 33 reads showing recombination evidence were categorized into 23 distinct events, with no single event carrying more than three reads.

While there is no definitive boundary between monoclonal and polyclonal samples, for simplicity we defined samples with SI greater than 0.25 (half of the upper limit of 0.5) as monoclonal and samples with SI less than 0.125 (half of the previous threshold) as polyclonal. Using these thresholds, 13 and 22 samples were classified as monoclonal and polyclonal, respectively, and 12 samples with SI values above 0.125 and below 0.25 were classified as in-between.

#### 2.1.3 Non-canonical V(D)J recombination events

There found a total 840 distinct V(D)J recombination events in the HPRC samples, with 546 (65%) being canonical, or complete V(D)J recombination events. In the remaining 294 (35%) distinct non-canonical events, 221 (26%) are partial (D-J only) recombination, and remaining 73 (9%) are other types of non-canonical recombination events.

In a canonical V(D)J recombination event, both the D-J recombination and V-D recombination utilize the same D gene. That is, the RSSs on both sides of one specific D gene are connected to other specific J and V genes via split alignments (Figure 2 **a**). A partial recombination event occurs when the recombination process is halted after the sequence between J and D genes is deleted. However, the sequence following the selected D gene is intact and remains in its germline form (Figure 2 **b**). Other types of non-canonical recombination events involve at least one additional recombination step after the D-J recombination. In these cases, two or more D genes may be involved (Figure 2 **c**). Additionally, we observed non-canonical recombination events that include sequence inversions or involve multiple J or V genes. Events like these often include non-coding regions of the IGH locus and are unlikely to produce a functional end-product. However, they can be confused with SVs in the germline genome and thus need to be discarded during downstream analyses.

**Figure 2:**
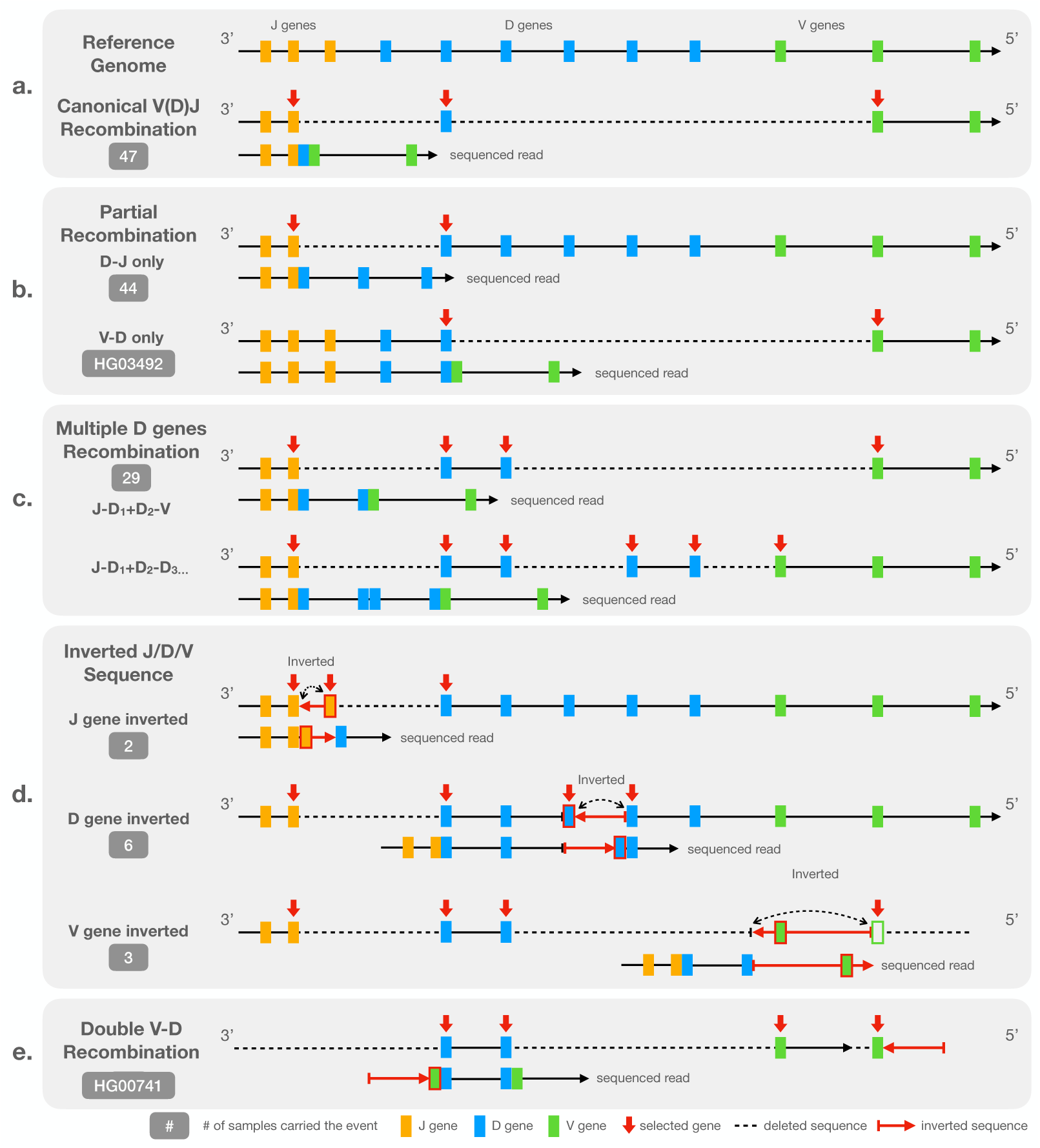
Diagram representation of **a**: a canonical V(D)J recombination event relative to the reference genome, **b**: partial (D-J only or V-D only) recombination event, **c**: non-canonical events involving multiple D genes, **d**: non-canonical events involving inverted V, D, or J sequences, and **e**: a rare non-canonical event involving two V-D recombinations. The “sequence read” shows the pattern we observed in the HiFi read while the above representation shows how is the read recombined from germline reference genome.

Figure 2 **b** to **e** illustrates the observed types of the non-canonical recombination events, which we also detail here:

##### Partial (”D-J only” or ”V-D only”) recombination

The ”D-J only” recombinations are the most prevalent non-canonical recombination event found in 44 out of the 47 samples.

Note that the number of partial recombination events we detected can be an overestimate because the beginning part of the sequenced read between partial recombination and “multiple-D-genes” events are identical. A “multiple-D-genes” read can be mistaken with a partial recombination if the read is not long enough to span the second recombination event. However, the distinction between partial recombination event and canonical recombination is evident due to the existence of the germline flanking sequence on the 5*^′^* end of the chosen D gene. The lengths of human D genes do not exceed 40 bp making them easy to span by HiFi reads.

Nearly all observed partial recombination events involve ”D-J only” recombination exclusively. However, an exception was noted in the sample HG03492 (Figure S5), where ”V-D only” recombination occurred independently. In this case, the sequence on the 3*^′^* end of the D gene split sites extend to the region between IGHJ and IGHD loci, indicating the absence of the D-J recombination in the event.

##### Multiple D genes

The second most common type of non-canonical recombination events involves more than one D gene and was detected in 29 out of 47 samples in HPRC.

In this scenario, V(D)J recombination uses the 5’ RSS of one D gene and the 3’ RSS of another D gene located downstream from the first one thus creating a virtual ”ultralong” (and not necessarily productive) D gene. In (18), this scenario was referred to as *RSS-skipping*. A simple example of RSS-skipping is illustrated on an IGV screenshot that shows alignments of HiFi reads from HG02557 (Figure S6). The red arrows highlight a read that is split-aligned into three segments, with two long deletions crossing (*IGHJ2*, *IGHD4-23*) and (*IGHD6-19*, *IGHV3-49*). Note that the *IGHV3-49*-end segment is outside the bounds of the screenshot. The recombination event utilizes the 3*^′^* RSS of *IGHD4-23* and the 5*^′^* RSS of *IGHD6-19* rather than the two RSSs of a single D gene. A more complex example of RSS-skipping involving three D genes –*IGHD5-18*, *IGHD4-17*, and *IGHD2-2*– is indicated with blue arrows in Figure S6.

##### Inverted J, D, or V genes

Figure 2 **d** shows how J, D, or V genes can be inverted in a recombined genome. This type of non-canonical event is rarer compared to the partial or multiple-D-genes event with 2, 6, and 3 samples carrying J, D, and V gene inversion events, respectively.

In instances where the J sequence is inverted, the sequence between the 5*^′^* RSSs of two consecutive J genes is cropped and inverted. The two J genes therefore appear connected on the 3*^′^* end of the inversion in the sequenced read. We showed an example of inverted J gene with individual HG02559 in Figure S7.

In the case of D genes, the sequences between the 3*^′^* RSSs of two consecutive D genes are cropped and inverted. As for the J gene, the two D genes are connected together at the 5*^′^* end of the inversion in the sequence read. An example of inverted D sequence with individual HG01978 is shown in Figure S8.

Finally in the case of V gene, the structure is different to the previous two instances. The inversion is observed between one D and one V gene. In a canonical V(D)J recombination event, the 5*^′^* end of the D gene is connected to 3*^′^* end of a V gene with some junctional diversity. However in the inverted V sequence cases, the 5*^′^* end of the D gene is connected to the inverted 3*^′^* flanking sequence of a V gene including the RSS of the V gene. An inverted V sequence example with individual HG00621 is shown in Figure S9.

The inverted sequence event can also combined with “multiple D genes” event. All the inverted D gene cases involved in multiple D genes by nature. The three inverted V gene cases are all combined with two different D genes. In one of the two cases where an inverted J sequence was observed, we also observed multiple D genes, combining the effects shown in Figures 2 c & d. In the other case, the read is not long enough to distinguish whether one or more D genes are involved.

##### Double V-D recombination

We observed this type of non-canonical recombination events only in one individual (HG00741). The event consists of one recombination event involving the 3*^′^* RSS and 5*^′^* RSS of two consecutive D genes (Figure 2 **e**). Both the D gene RSSs are adjacent to a V gene, creating an amalgam of two connected V-D recombination sequences. No J gene is involved in the event. We hypothesize that the J gene was deleted during the recombination event. An IGV screenshot detailing the evidence for this event – which spans multiple reads – is shown in Figure S10.

Non-canonical V(D)J recombination events are common across the samples, accounting for around 35% of all recombination events. They include long range rearrangements such as deletions and inversions between RSSs of the V, D, and J genes and are arranged in ways different from canonical V(D)J recombination. Understanding the mechanisms behind these events is essential for differentiating reads carrying somatic events from reads carrying germline SVs in the IGH locus.

### 2.2 IGLoo --asm measures the somatic effect on the HPRC assemblies

For LCL datasets, V(D)J recombination impedes the assembler’s ability to accurately assemble the germline genome sequence. For example, recombination depletes evidence from the the germline sequence between the V and J genes. Polyclonality leads to emergence of reads originating from somatically rearranged haplotypes, which might violate the assembler’s expectation of a strictly diploid and nearly uniformly covered genome. Consistent with this, we observed that the HPRC assemblies were not contiguous in the IGH locus, with IGH genes have been lost during assembly.

The `IGLoo --asm` module is designed to profile the IGH locus in genome assemblies, locate breakpoints in the contigs with respect to the germline IGH genes, and identify IGH genes missing from the assembly. `IGLoo --asm` first calls gAIRR-suite (25) to annotate the IGH genes on the assembly. It identifies all contigs overlapping IGH genes and filters out the contigs containing only IGH genes with more than 15 mismatches. These are likely to contain “orphon” genes, which have homology to IGH genes but are located outside the main IGH locus (Method section 4.2). Finally, `IGLoo --asm` compares the filtered contigs to the germline IGH locus to situate breakpoints with respect to the IGH genes.

We used `IGLoo --asm` to profile all of the genome assemblies from the HPRC project. Figure 3 **a** shows the number of contigs for the 94 HPRC haplotype assemblies (two assemblies per individual). 21 (22%) of the haplotypes have a single contig covering all or part of the IGH locus. 28 (30%) of the haplotypes have the IGH locus spilt into two contigs, and 45 (48%) haplotypes have the locus split into three or more contigs. In cases where the IGH locus was covered by a single contig, the assembly sometimes suffered from missing genes due to V(D)J recombination events or other assembly issues. Figure 3 **b** shows the total number of V genes (without pseudogenes), D genes, and J genes covered by the longest contig for each haplotype. According to IMGT (24), there are 95 such genes in the IGH locus, but the number varies due to germline variation. Because most of the insertions in the IGH locus are annotated within the 95 genes, it is more common for a haplotype to carry fewer genes than to carry more.

**Figure 3:**
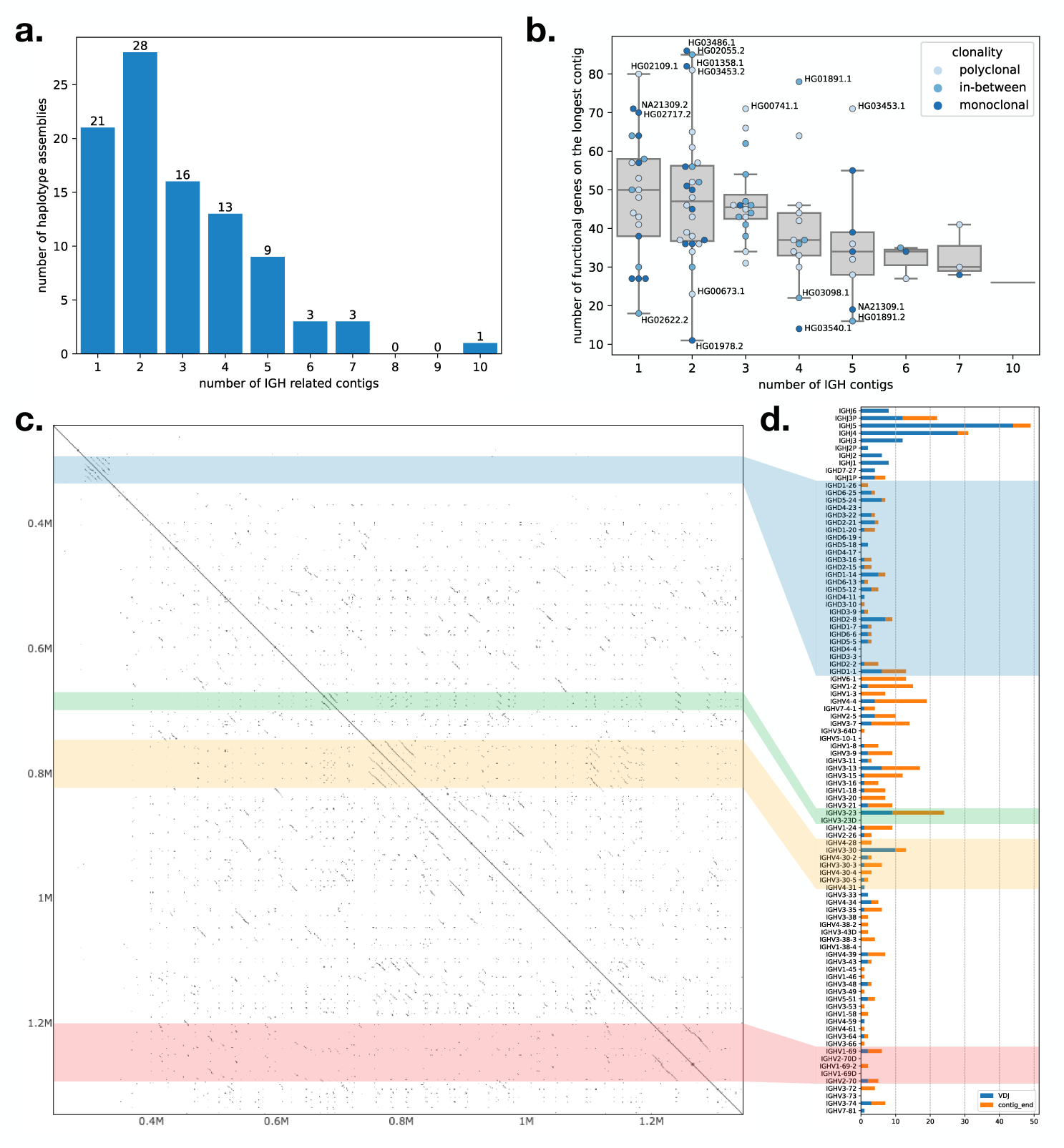
**a**. the accumulated bar plot of the number of contigs in one haplotype assembly of HPRC. **b**. the number of functional IGH genes covered by the longest contig in each haplotype assembly. Dots represent haplotype assemblies, and the hues of the dots represent the clonality of the cell line according to Simpson’s index and our definitions of monoclonal and polyclonal samples in section 2.1.2. **c**: self-aligned dot plot of the sample NA19240 haplotype 1 assembled by `IGLoo` with ModDotPlot (26). **d**: the accumulated breakpoints of the 94 HPRC haplotypes. Breakpoints are classified as with V(D)J recombination evidence or not, e.g. including the disjoint, overlapping, and duplication cases in Figure S11. The colored strips across the **c** and **d** panels show the relative position of the repetitive regions in the dot plot and bar plot.

A joint analysis of sample clonality and assembly continuity did not reveal clear relations between these two factors (Figure 3 **b**). For example, of the 21 haplotypes on which the IGH locus assembled into a single contig, around half carried fewer than 50 IGH genes (Figure 3 **b**). On the other hand, some of the assembled haplotypes that were broken into two contigs, e.g. HG03486’s paternal and HG02055’s maternal haplotypes, each carried more than 80 IGH genes.

Another challenge to correct assembly of the IGH locus is the presence of repeats. We focused on four common repetitive regions in the part of IGH locus spanning V, D, J genes illustrated in a self dot plot for NA19240’s paternal haplotype (Figure 3 **c**). The dot plot showed the repetitive regions in the self alignment of the complete NA19240 paternal IGH locus we reassembled in section 2.3. The four regions with parallel diagonal lines are the common repetitive regions on the IGH locus: the IGHD gene locus (the blue block), the duplication of the *IGHV3-23* (the green block), a nearly tandem repeat containing *IGHV3-30*-like and *IGHV4-30*-like genes (the yellow block), and the locus containing copies of IGHV1-69 genes (the red block). Previous studies show that these loci represent hotspots for SVs (3), and some haplotypes in the HPRC population are also characterized by different number of copies of *IGHV3-23*, *IGHV1-69*, or *IGHV3-30*-like genes.

To analyze the impact of repetitive regions on assembly continuity, we examined the IGH locus assembly for breakpoints and checked whether their distribution overlaps with the repetitive regions. We defined a breakpoint as either the discontinuity point between two contigs or the position where the assembly transitions from germline sequence to somatic haplotype. The first type of breakpoints, which we called “contig ends”, occur at the ends of contigs that do not span the entire IGH locus, leaving contig ends within the locus. Two adjacent contigs can be disjoint, overlap each other, or one can be contained in the other (as shown in the top three cases in Figure S11). In all these cases, the breakpoints are defined as the ends of the contigs. The causes of these breakpoints are discussed below.

The second type of breakpoints, which we called “V(D)J breakpoints”, are the points in the assembly that show evidence of V(D)J recombination. We detected these “V(D)J breakpoint” by searching for contigs connecting from V to J, V to D, or D to J genes (Figure S11 bottom). Although the “V(D)J breakpoint” technically does not affect assembly continuity, they can contribute to genes being missing from the assembly. We marked the closest J, D, or non-pseudo V genes to the breakpoints in the 94 HPRC assemblies, summarizing them in a bar chart (Figure 3 **d**). The colored strips indicate which repeat regions in the dot plot (Figure 3 **c**) correspond to these bars. Since all the D genes except *IGHD7-27* are included in the D gene locus repeat, the blue transparent block roughly divided the J/D/V genes in the Figure 3 **d**. All the genes below the block are the V genes, and the genes above the blue transparent block except *IGHD7-27* are J genes.

In Figure 3 **d**, the “V(D)J breakpoints” are enriched in the J genes, consistent with the fact that there are fewer J genes. On the other hand, “contig end” breakpoints seem not to be enriched in repetitive regions, but instead are enriched in the area between *IGHV6-1* to *IGHV3-23*, which has only a small overlap with the *IGHV3-23* repeat region. We concluded that the fragmentation of the IGH locus is mainly due to V(D)J recombinations, and not to repeat regions. Note that the cause of “contig end” breakpoints is not as evident as that of “V(D)J breakpoints”. “Contig end” breakpoints can be caused by presence of somatic haplotypes or a a drop of read coverage for germline sequences, which are depleted during recombination. This read depletion is expected to be especially severe in areas near the D gene locus, consistent with the “contig end” breakpoints we observe.

### 2.3 IGLoo --ReAsm can improve the IGH assemblies

Assemblies derived from LCLs can be fragmented or can contain other flaws. The `IGLoo --ReAsm` module improves the accuracy of the germline assembly obtained from LCLs. This module deploys two distinct reassembly methods, one that uses the hifiasm *de novo* assembler (16) to assemble the backbone of the IGH locus. And then the backbone is further improved by the reference-guided assembler MaSuRCA (17) (Figure 4 **a**).

**Figure 4:**
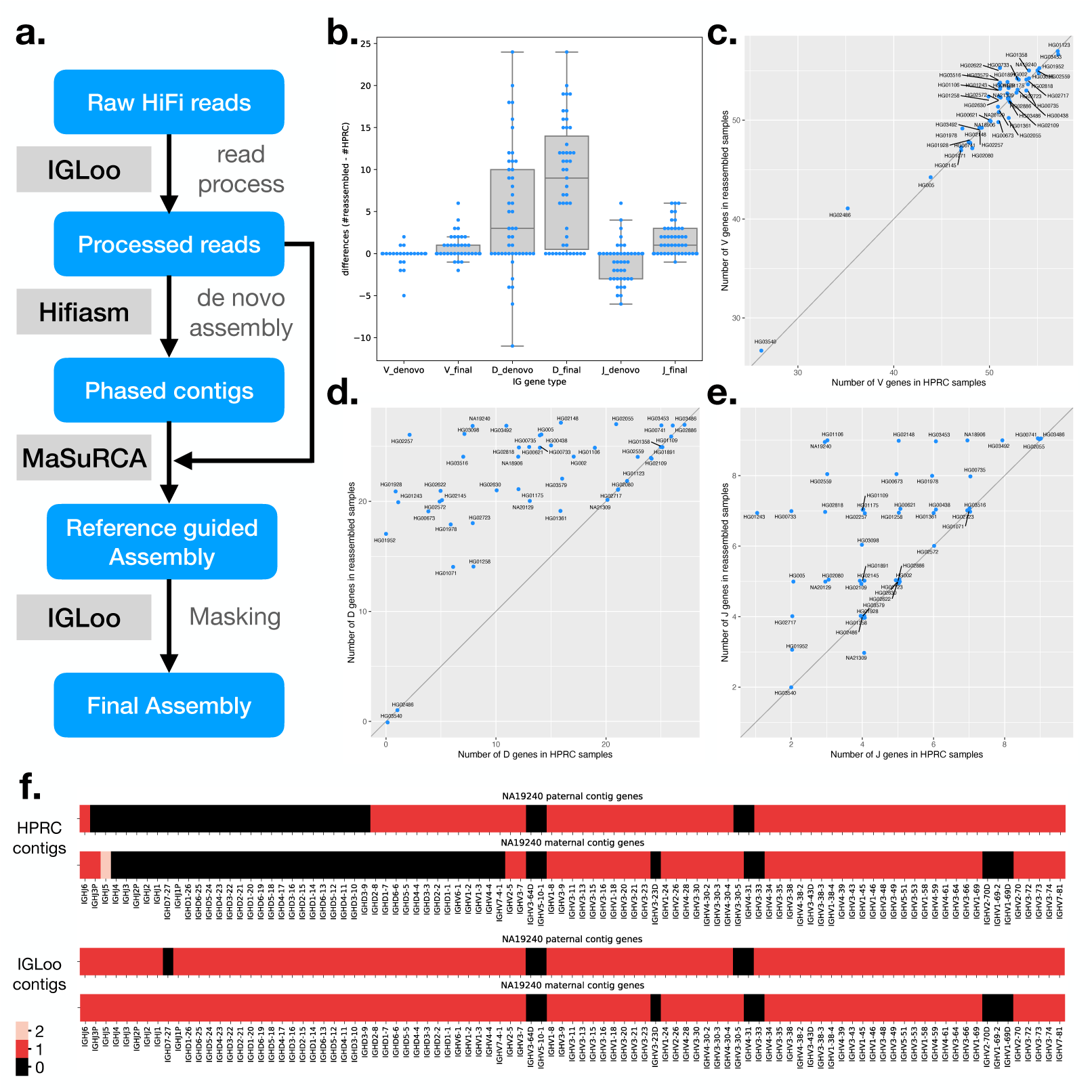
a. Simplified `IGLoo --ReAsm` assembly pipeline. The full assembly pipeline is in section 4.3. **b.** The distribution of gene number differences between *de novo* assembly contigs and HPRC assemblies, and the differences between final assemblies and HPRC assemblies. Each dot represent one sample. Notice that there are more dots crowded in the 0 entries, which exceed the width of the box. **c.** The V gene number differences of HPRC assemblies and reassembled IGH locus. **d.** The D gene number differences. **e.** The J gene number differences. **f.** the gene number in its relative position on the sample NA19240 before and after `IGLoo --ReAsm`. All D and J genes are analyzed, but only functional or ORF V genes are analyzed for simplicity.

Two main challenges of the LCL data are the somatic haplotypes and the low read depth around the J and D gene loci. The V(D)J recombination events in the data set create non-germline haplotypes, which the assembler cannot automatically disregard in favor of the germline haplotype. Low depth in the region between J and D genes causes fragmentation and loss of IGH genes.

To tackle the somatic haplotype problem, `IGLoo --ReAsm` preprocesses the raw HiFi reads before reassembling them with hifiasm. Based on the analysis of `IGLoo --read`, `IGLoo --ReAsm` collects all the reads showing evidence of V(D)J recombination breakpoint, indicated by split-alignments on two sides of IG gene RSSs. These reads are then split at the breakpoint positions. To mitigate the effect of reference bias, `IGLoo --ReAsm` selects the best mapping result from the three reference genomes used by `IGLoo --read` (Method Section 4.1). By doing the splitting, the somatic haplotypes are broken. On the contrary, the germline haplotypes remain intact in the read pool.

`IGLoo --ReAsm` also performs read enrichment to counter the read depletion in the region. Enrichment is achieved by creating artificial copies of reads mapping to the region between J and D genes, approximately doubling the read depth in the region and giving the assembler a stronger basis for assembling the region without fragmenting the contigs and losing D genes.

#### 2.3.1 *De novo* assembly results

To produce an initial version of the improved germline assembly, `IGLoo --ReAsm` runs hifiasm with the trio binning method. Rather than use the full set of Illumina short reads from the individual’s parents, we extract and use only the parental reads mapping to the IGH locus. Since the parental data also originate from LCLs, it can suffer from the presence of somatic haplotypes and low read depth in the IGH locus thus making the phasing information less reliable compared to other regions of the genome. Because somatic haplotypes are hard to identify using Illumina short reads, we do not attempt to remove reads containing V(D)J recombination events from the parental datasets. The assembled contigs are later used as the backbone for next stage assembly and polishing.

We use the `IGLoo --asm` to evaluate our reassembled genomes. Figure 4 **b** shows the difference in the number of IGH genes between HPRC assemblies and the *de novo* assembled contigs from processed reads. Boxes marked “ denovo” show the difference in count of the V, D, and J genes between HPRC and *de novo* assemblies on processed reads. For most individuals, the number of V genes stayed roughly equal, with the average number of differences being *−*0.15. The number of D genes increased, with the average change being +4.87 per individual. However, the number of J genes decreased by *−*0.94 on average.

The increase in the number of D genes can be explained by the preprocessing of the HiFi reads, which removes the somatic haplotype and increases the read depth in D gene locus. The decrease in the number of J genes is more complicated. Since most recombination events involve a J gene, raw reads covering J genes are likely to span a split site. As a result, preprocessed reads covering J genes are often clipped near a J gene. This clipping, combined with the possibly presence of SHMs and class switch recombination events occurring at the 3’ end of the J gene region, can cause fragmentation in the J gene locus. On the other hand, assembling from raw reads can generate some somatic haplotype, in which a few J genes can still be preserved.

#### 2.3.2 Reference guided assembly

To recover the lost J genes in the *de novo* assembly stage, and to polish the assembly result, `IGLoo --ReAsm` uses the reference-guided assembler MaSuRCA (17). First, `IGLoo --ReAsm` uses the information in *de novo* assembled contigs to generate a personalized reference genome for each of the individual’s haplotypes. The contigs, personalized reference, and processed reads are then fed to the MaSuRCA pipeline for scaffolding, gap filling, and polishing. Finally, `IGLoo --ReAsm` masks portions of the assembly that lack read support, since these portions come from the reference provided to MaSuRCA, rather than from the input reads.

An SV allele is defined as a large haplotype that differs from the reference genome. `IGLoo --ReAsm` is able to detect the prevalent SV allele type from the *de novo* assembled contigs. Sometimes, the allele type of a contig can contradict that of the other contig in the same paternal or maternal group. The contradiction may come from mis-categorizing of a contig from the other parental group due to the issue stated in section 2.3.1. `IGLoo --ReAsm` detected the SV haplotype and attempted to reconcile allele types by swapping conflicting contigs between paternal and maternal groups. Three samples –HG01891, HG03453, and NA18906– could not be reconciled through contig swapping. We addressed these three samples manually. After the reconciliation, a personalized reference is generated. `IGLoo --ReAsm` used the backbone of T2T-CHM13, and stitched different haplotype of the SV according to the information provided by *de novo* contigs. (Method section 4.3.4)

After the reference guided assembly, `IGLoo --ReAsm` realigned the HiFi reads back to the MaSuRCA output and masked out regions without any support from the read alignments or *de novo* assembled contigs. These are regions that likely originating from the guide reference rather than from the input reads.

We again used `IGLoo --asm` to evaluate the final masked assemblies. Figure 4 **c, d**, and **e** show the differences in V, D, and J gene counts between HPRC assemblies and the final masked `IGLoo` assemblies. Each dot represents an individual, and most dots are located on the diagonal line or in the upper left triangle, showing that `IGLoo` usually recovered an equal or greater number of IGH genes compared to the original HPRC assemblies. Figure 4 **b** shows the gene number differences from HPRC assemblies. The box marked with “ denovo” are the numbers of *de novo* assembled contigs, and the numbers of the final assemblies are shown in the boxes marked with “ final”. In general, *de novo* assembly recovered some D genes while losing some J genes. On the other hand, the reference guided method recovered the missing J genes from *de novo* assembly, and also recovered more D genes. Note that in Figure 4 **b**, some of the 0-difference dots are not shown because they exceed the width of the box. Out of the 47 samples, 17 had an increased number of V genes and 4 had a decreased number. 35 samples had an increase in *D* genes. 29 had an increased number of J genes and one has a decreased number. The average V, D and J gene differences were 0.62, 8.51, and 1.64 respectively.

Figure 4 **f** illustrates one example NA19240 of HPRC assemblies alongside the final `IGLoo` masked assemblies. The genes on the contigs are arranged by the IGH locus. Both haplotypes of the HPRC assembles exhibited V(D)J recombination events, whereas the `IGLoo --ReAsm` assembly recovered more continuous haplotypes. It should be noted that the maternal haplotype reconstructed by `IGLoo --ReAsm` may not accurately represent the germline haplotype. Upon manual inspection, the read sequences mapped to the D gene locus show haploid evidence, and the genotype resembles the paternal data (Figure S12). This may indicate that the D gene information originated from the paternal haplotype only.

#### 2.3.3 Comparison with IGenotyper

IGenotyper (14) is another method for profiling and assembling the IGH locus. We applied IGenotyper to the HPRC dataset in order to compare it to `IGLoo`. For efficiency, we used the same inputs as were provided to the `IGLoo --ReAsm` pipeline, where the reads have already been aligned (by HPRC) and filtered (by `IGLoo`) to the IG loci.

We ran IGenotyper on the 47 HPRC samples without conducting any manual curation as described in (14), we used the output file `igh contigs.fasta` from IGenotyper assembly step. We employed `IGLoo --asm` to evaluate the IGenotyper assemblies and compared them to the `IGLoo --ReAsm` results in the same manner as the comparison with HPRC assemblies. Figure S13 **a** illustrates the differences in IGH gene numbers between the two methods. The distribution of the differences resembles the comparison between HPRC assemblies and `IGLoo --ReAsm`. Specifically, while the number of IGHV genes remains comparable, `IGLoo --ReAsm` exhibits a significant increase in IGHD and IGHJ genes. On average, the V gene difference is 1.13, D gene difference is 11.70, and the J gene difference is 3.89. Figure S13 **b** shows the total IGH gene numbers in the samples between the two methods. All 47 samples either favor `IGLoo --ReAsm` or show no difference in the gene numbers. This is as expected, since `IGLoo --ReAsm` additionally removes somatic haplotypes before assembly.

## 3 Discussion

We presented the `IGLoo` software tool, which implements several new methods for elucidating both the germline and somatic natures of the IGH locus when sequenced using HiFi reads from an LCL. `IGLoo` fills a gap in the current landscape of tools for *de novo* assembly of HiFi data, which treat data derived from LCLs as though it represented a pure germline sequence.

We used `IGLoo` to study the non-canonical V(D)J recombination events in the HPRC assemblies, finding that non-canonical events made up approximately 35% of all recombination events in the cell line. The length of HiFi reads facilitates analysis of somatic events involving IG genes inversions and recombination of multiple D genes. Understanding these recombination events aids in profiling the clonality of the LCLs in HPRC and allows one to distinguish variations driven by V(D)J recombination from germline SVs. We focused on the IGH locus and did not analyze off-target recombinations (recombinations outside of IG loci), though we hypothesize that cryptic RSSs might drive additional rearrangements and thus complicate the inference of germline SVs. Further, we showed specific ways in which somatic recombination events led to assemblies that were either fragmented or failed to represent the germline sequence. Given our understanding of the V(D)J recombination mechanism and the population diversity in the IGH locus, we contend that `IGLoo` will be an important tool to improve personalized assemblies, especially since it recovers IGH genes that would be missed by standard assembly tools.

We found that *de novo* assembly of preprocessed reads improved the representation of the D gene locus while also resulting in some missing J genes. Although the number of D genes gained is greater than the number of J genes lost, there remains a trade-off between the two. Reference-guided assembly appears to recover both D and J genes without this compromise. However, we observed that the reference-guided assembler MaSuRCA occasionally favors the reference genome over *de novo* assembled contigs, especially when structural differences exist between them. Therefore, it’s crucial to construct a personalized reference genome that closely resembles the sample. `IGLoo` constructs this personalized reference genome based on common SVs reported by (3)). However, we also noted SVs that deviate from these common ones. In such cases, `IGLoo` cannot generate a reference genome that includes these special SV types, presenting a challenge for MaSuRCA to successfully assemble the region. In the future, as more IG SVs are profiled and cataloged, `IGLoo` will be able to generate a more accurate personalized reference genome.

In the future, it will be also important to study applicability of the method to immunoglobulin light chain loci and examine other somatic recombination events within the IGH locus, e.g. class switch recombination events involving deletions between the J and C genes, which we did not address. However, more work and customization will be needed to handle the particulars of these loci and events.

The `IGLoo` reassembly pipeline serves as a workaround for assemblers that are not designed to handle somatically recombined haplotypes. Hifiasm was designed to assemble phased assemblies based on trio data. However, it does not take into account the fact that parental data can undergo V(D)J recombination, resulting in unbalanced coverage in the IGH locus. The substantial read depletion in the IGH locus also poses a significant challenge to hifiasm’s *de novo* assembly approach. While MaSuRCA handles the read depletion issue more effectively in the dataset, it lacks the ability to provide a phased assembly. Transferring hifiasm’s phased results to MaSuRCA partially addresses this issue. Nevertheless, there may still be instances of phasing errors or over-correction by MaSuRCA. A specialized assembler tailored for the IG loci, which considers parental data and addresses the read depletion issue within the locus, could potentially yield more accurate IGH assembly results.

The `IGLoo` modules are specifically designed for analyzing HiFi datasets and conducting assembly based on HiFi reads. However, the `IGLoo --read` module can potentially be adapted to handle pair-end short read data. By examining the aligned positions of both ends of a read pair, a similar analysis to split-alignment of HiFi reads can be applied.

On the other hand, detecting non-canonical recombination events would not as straight-forward as with HiFi reads. Full non-canonical recombination events likely cannot be captured by a single read pair, requiring some form of inference similar to those used in transcriptome analysis. Another potential challenge for short reads is the compact nature of the J genes in the IGH locus as the distance between two consecutive J genes does not exceed 400 bp. If the non-sequenced insert of the pair-end read falls within the J locus, determining which J gene contributes to the recombination event can be ambiguous and may necessitate inference techniques.

Profiling and assembling IG loci based on non-LCL sources (e.g. PBMCs) could be considered a less challenging approach for constructing references for IG loci. However, since LCLs continue to be widely used and abundant in existing studies, `IGLoo` offers valuable insights for researchers, aiding in the comprehension of IGH locus diversity. `IGLoo` reconstructs the IGH locus, which is often overlooked in standard assembly pipelines. Although the resulting assembly may not be perfect, it represents a significant step closer to obtaining the best possible results from cell line data.

## 4 Methods

### 4.1 Profiling V(D)J recombination events using HiFi reads

#### Split alignments and deletions

`IGLoo --read` accepts a HiFi read file (FASTA/FASTQ) or a read alignment file (BAM/CRAM) as input. When processing alignment file, `IGLoo --read` extracts the read aligned to IGH locus with `samtools view` and the BED file specified the IGH locus on the reference genome. When processing a read file, `IGLoo --read` uses `minimap2` to align the reads, then extracts the IGH-locus alignments.

`IGLoo --read` then analyzes the read alignments, starting from the leftmost J gene to the rightmost D gene. Reads providing evidence of a V(D)J recombination event will typically align in a split fashion across two or more locations on the reference genome. We refer to the distinct portions of the read that align contiguously as “segments.” `IGLoo --read` identifies split alignments and counts such an alignment as “meaningful” if at least one of the junctions in the split alignment is near the RSS of any V, D, or J gene. Our analysis on HPRC data indicates that the majority of the split sites (94%) fall within 50 bp of an RSS, so we defined the split alignments as sufficiently close if they are within 50 bp apart from RSSs of any IGH genes (Figure S1).

Additionally, `IGLoo --read` examines the deletions within each “segment”. In cases of non-canonical V(D)J recombination events between D genes, two consecutive D genes may be close enough that the aligner represents the event as a deletion rather than a split alignment. `IGLoo --read` analyzes each alignment’s CIGAR string, identifying a long deletion between two D genes as indicating a recombination event. Due to the repetitive nature of the D gene locus, the aligner may make different decisions for different reads about where exactly to place the deletion. Hence, the split sites of the deletions may not necessarily be close to an RSS. Therefore, for recombination events between two D genes, we set a loose requirement that the length of deletion be within 50 of the distance between the two relevant D genes.

#### Multiple references

To counter reference bias in the IGH locus, we use three reference genomes: GRCh37, GRCh38, and T2T-CHM13. Each reference has some IGH genes not shared by the others. For example, the IGH genes *IGHV1-8* and *IGHV3-9* are common in the population but exist only in GRCh37 and not the others. GRCh38 carries the gene locus from *IGHV4-30-2* to *IGHV4-31*, not present in the other two. Only T2T-CHM13 carries the duplication of gene locus from *IGHV2-70D* to *IGHV1-69D* (Figure S14).

Each extracted read is aligned to all three reference genomes. Of the three choices, `IGLoo --read` picks the one that best fits the reference genome and its annotation. For instance, an alignment on one reference genome is preferred if all of its split sites are close to RSSs in that reference.

`IGLoo --read` then classifies each “split” alignment to the kind of event that it supports. An alignment is classified as supporting a “canonical V(D)J recombination event” if it has a single split, with one segment aligned to a J gene and the other segment aligned to a V gene, with split sites situated near the RSSs of the respective genes. Note that for V(D)J recombination events, the portion of the D gene present in the read is too short to induce the aligner to make two separate splits on either side of the D gene. As a result, a canonical event will join one J gene to one V gene, with the D gene sequence appearing on one of the segments, or possibly soft clipped in the alignment. An alignment is classified as supporting a “non-canonical” event in two scenarios. Firstly, if an alignment is split once, with one segment aligned to a J gene and the other aligned to a D gene, it indicates evidence of a J-D recombination event. Alternatively, if the read is split into multiple segments, it suggests a “non-canonical” event involves multiple deletions.

### 4.2 Analyzing the personal assemblies

To analyze the quality of the IGH locus in a personalized assembly, `IGLoo --asm` first runs gAIRR-annotate (25) on the assembly. gAIRR-annotate uses BWA MEM (27) to align documented IGH genes/alleles from the IMGT database to the assembly, reporting the closest genes/alleles and their positions on the assembly. `IGLoo --asm` then evaluates the assembly based on the distribution of IGH genes on each contig. With the information, `IGLoo --asm` reports the contiguity of the IGH locus, identifies missing IGH genes, and pinpoints the breakpoints in the assembly for assembly quality assessment.

Outside of the main IGH locus on chromosome 14, there are also many “orphon” genes distributed across the genome. These are possible pseudogenes that are homologous to the main IGH locus. Though they share sequence similarity to IGH genes, our focus is solely on the main IGH locus, and we do not aim to improve assembly quality at orphon sites. We therefore filtered out orphons as well as any IGH genes with more than 15 mismatches compared to the documented genes, which are likely non-functional genes. `IGLoo --asm` then filtered out the contigs containing only these genes and retaining the contigs with functional IGH genes in the downstream analysis.

`IGLoo --asm` provides a report on the IGH gene structure, as illustrated in Figure 4 **f**. However, it’s worth noting that the output from gAIRR-annotate, which is solely based on sequence, does not distinguish between duplicated gene pairs such as *IGHV3-23* and *IGHV3-23D*, *IGHV1-69* and *IGHV1-69D*, or *IGHV2-70* and *IGHV2-70D*. To improve interpretability, `IGLoo --asm` correctly assigns duplicated genes to their respective names. The process involves first tallying the occurrences of identical genes, then assigning duplicated genes with “D” in their name only if multiple instances are detected within the contig. While there are other types of duplicated genes in the IGH locus, they can be easily differentiated by considering their neighboring genes.

After the filtering and gene assignment steps, `IGLoo --read` counts the number of contigs and finds the breakpoints in the assembly. We classified the breakpoints into two main types: “V(D)J breakpoints” or “contig end”. V(D)J breakpoints do not cause the contig to become fragmented, but do cause the contig to contain the recombined sequence. We detect the V(D)J breakpoints by scanning the IGH genes on each contig. If two IGH genes from different loci, i.e. V and J loci or D and J loci, are located next to each other within 10 kbp, the connection is considered a V(D)J breakpoint. The contig end breakpoints correspond to situations where two contigs cannot be connected into one, fragmenting the assembly. The relation between the two contigs can further be categorized into “disjoint”, “overlap”, or “duplication” as depicted in Figure S11. To collect the contig end breakpoints, we first sort the contigs according to their relative position on IGH locus, then check and compare the IGH genes on the boundary of the nearby contigs.

### 4.3 Reassembly to improve germline assembly of IGH locus

`IGLoo --ReAsm` performs a series of steps to reassemble the genome to better capture the original germline sequence, rather than the somatically recombined sequence. The workflow is shown in Figure 5. `IGLoo --ReAsm` first preprocesses the HiFi reads, filtering and duplicating some reads in order to shift the read evidence away from areas with V(D)J recombination and toward areas that clearly represent the germline. Preprocessed reads are then given to hifiasm (16), along with k-mers counts from sample’s parents. Hifiasm then uses trio binning (28) to generate a phased personal assembly. `IGLoo --ReAsm` then checks the contigs to infer the allele type of some known SVs in the paternal and maternal haplotypes. Based on these, `IGLoo --ReAsm` generates both paternal and maternal personalized reference haplotypes. The personalized haplotypes, combined with the hifiasm contigs and preprocessed reads, are then given to the reference guided assembler MaSuRCA to generate high quality IGH assembly (17). Finally, `IGLoo --ReAsm` masks out the regions in the high-quality assembly to eliminate any regions whose sequence came only from the reference sequence provided to MaSuRCA, and which were not influenced by the HiFi read sequences.

**Figure 5:**
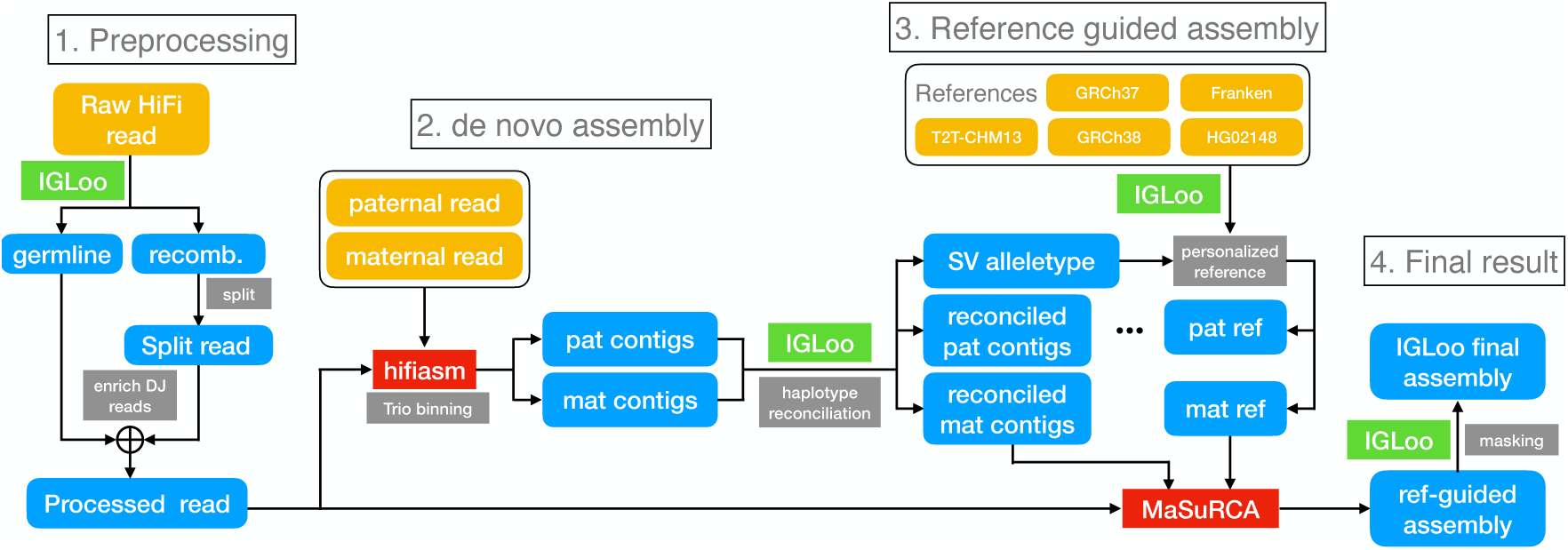
The full IGLoo --ReAsm assembly pipeline.

#### 4.3.1 Preprocessing the HiFi reads

This module consists of two steps. The first step filters out read-level evidence originating from recombined portions of the somatic haplotype. Whether a read carries evidence for a somatic event was determined earlier in the `IGLoo --read` analysis of the split-mapped reads. Consequently, we partitioned the reads into two or more segments based on how many segments their alignment on the reference genome. This effectively eliminates evidence from the somatic haplotype while retaining the rest of the read sequence. Note that `IGLoo --read` does not consider the non-reference connection between two V genes to be somatic V(D)J recombination because this could be evidence of SVs in the sample. Although somatic hypermutations can be present in the reads, `IGLoo --ReAsm` does not attempt to correct these as they generally will not affect the structure of the assembly. Correction of somatic hypermutations can also falsely remove real polymorphisms in the sample.

The second step performs selective enrichment of reads carrying germline evidence in key loci. Specifically, any read segments mapping to the J gene locus, D gene locus, or the region between J and D genes, are duplicated, yielding a second copy of each. This roughly doubles the depth of the germline sequence representation in the region, which often suffers from read depletion due to the V(D)J recombination events in the cell line. The enrichment step increases the chance that the assembler favors the germline sequences in its reconstruction.

#### 4.3.2 Hifiasm draft assembly

The preprocessed reads are *de novo* assembled with hifiasm (Figure 5 **2.**) In this version of `IGLoo --ReAsm`, parental short reads are required for the trio-binning mode of hifiasm to improve the contiguity of the assembly. When parental data are not available, users can use hifiasm’s standard mode, treating the primary assembly and secondary assembly as the “paternal” and “maternal” haplotype assemblies for the following stages.

The phased contigs assembled by hifiasm are then used to infer the personal IGH reference (Section 4.3.4) and serve as input scaffolding materials of MaSuRCA (Section 4.3.5)

#### 4.3.3 SV allele type and contig reconcile

We genotype the common SVs in the contigs of each parental assembly. We considered both the IMGT database and the findings of Rodriguez et al (3) and constructed the following list of common SVs:

1. deletion from *IGHD2-8* to *IGHD3-3*.
2. deletion of *IGHV7-4-1*.
3. complex haplotype of *IGHV3-64D* and *IGHV5-10-1*, or *IGHV1-8* and *IGHV3-9*.
4. deletion of *IGHV3-23D*.
5. deletion from *IGHV4-30-2* to *IGHV3-33*.
6. deletion from *IGHV4-38-2* to *IGHV1-38-4*.
7. deletion from *IGHV2-70D* to *IGHV1-69D*

We type the SVs by first calling gAIRR-annotate to collect the IGH genes on the contig. If one contig carries the genes neighboring the two ends of a deletion, but not the genes in the deletion, the contig is considered to support that deletion. On the other hand, if the contig carries the genes inside the deletion region, the contig does not support the deletion. If a contig does not cover the region, the allele type is “unknown” for that SV in the contig. Previous work used the term “complex” to describe an SV involving two haplotypes, one carrying *IGHV3-64D* and *IGHV5-10-1* and the other carrying *IGHV1-8* and *IGHV3-9*. We genotype this SV by searching for genes that are unique to one of the two haplotypes. If the contig does not carry any of the four unique genes, or carries unique genes from both haplotypes, then the allele type is “unknown”.

The `IGLoo --ReAsm` checks if there are SV allele type conflicts within paternal contigs or maternal contigs. If there are conflicted contigs, `IGLoo --ReAsm` tries to move the conflicted contig to another parental group. If the conflict can be simply reconciled by moving the conflicted contigs around, `IGLoo --ReAsm` report reconciliation successful and move to the next stage. Otherwise `IGLoo --ReAsm` report reconciliation fail and request manual reconciliation.

#### 4.3.4 Personalized reference genome

The personalized reference genome is built based on five assemblies. We use T2T-CHM13 as the backbone of the reference, and the sequence of the other genomes to form the alternate allele type of the SVs. Figure 6 shows the choices of the assemblies in the 7 common SVs stated in section 4.3.3. For the deletion SVs, i.e. the SV 1, 2, 4, 5, 6, 7, the solid lines show the no-deletion allele type sequence, and the dashed lines show the sequence with the deletion. The SV 3 is the complex SV that each haplotype got its unique IGH genes. We chose the sequence of T2T-CHM13 and GRCh38 for the two haplotypes. Since all the references lack a deletion in D locus, i.e. the first SV, we use the *de novo* assembled sequence of HG02148’s paternal haplotype as the reference. For the same reason, because all the three reference genomes’ SV 4 and 6 are deletions, we use the sequence of a custom linear IGH “Franken” reference genome from (3) for the no-deletion allele type sequence.

**Figure 6:**
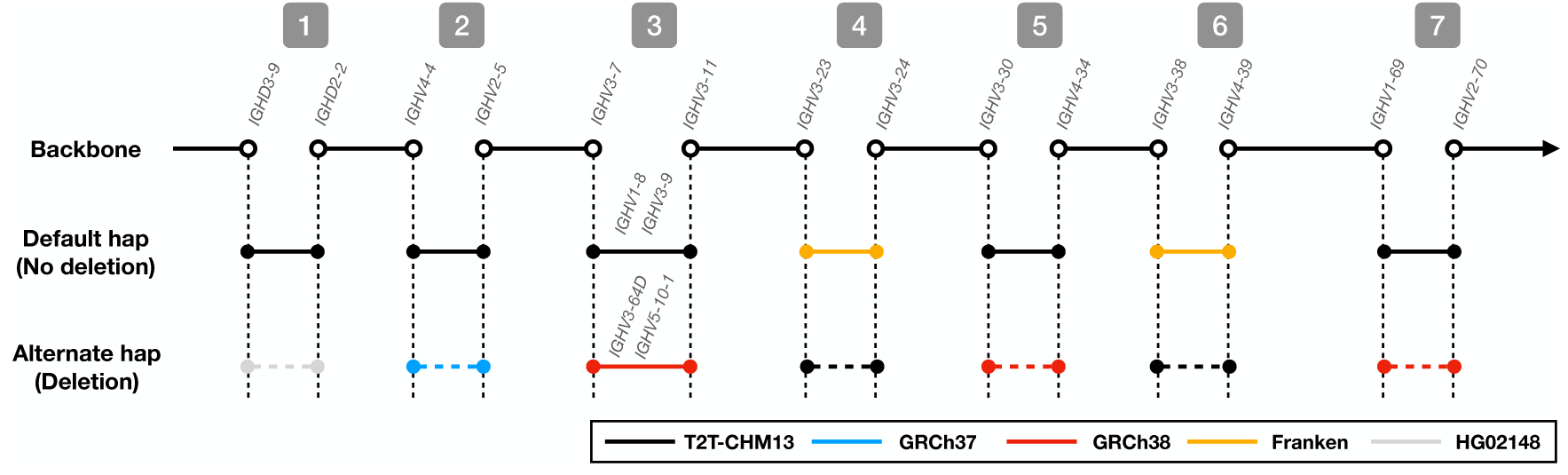
The sequence of choices made when building a personalized reference genome according to SV allele type.

We chose the nearby IGH genes as the anchor points for the sequences. The solid and hollow circles show how certain genes are chosen for inclusion in the sequence. Using the SV allele types from the previous stage, a personalized reference genome for each parental haplotype is created.

#### 4.3.5 Reference-guided assembly with MaSuRCA

We use the personalized reference as the guided reference of MaSuRCA, the *de novo* assembled contigs as the scaffolding materials. MaSuRCA first patches the contigs with the reference genome as the draft assembly. This causes gaps between contigs to be filled with sequence from the reference. We also aligned the processed reads to the two draft assemblies. Most reads will map approximately equally well to both draft assemblies, but a few reads map only to one assembly. The commonly mapped reads and uniquely mapped reads are collected to polish their respective assembly.

#### 4.3.6 Masking false-positive regions

Since the output contains some “patched” sequences taken from the reference, we filtered out regions without support from either contigs or reads. MaSuRCA uses the “capitalized nucleotide” to represent region in the assembly support by contigs or polished by the reads. We first discard MaSuRCA contigs not supported by any *de novo* assembled contigs. Then for the remaining MaSuRCA contigs, we align the processed reads to them. We use `samtools depth -a -g 0x100 -J` to check which regions are covered by read alignments. For regions without any coverage, we mask the region with “N”s.

### 4.4 Experiments perform on HPRC samples

We applied the `IGLoo` analysis pipeline to the 47 individuals of the HPRC year 1 release. Though not in our released software, we used GNU parallel (29) in our experiments.

In the Hifiasm trio-binning stage of `IGLoo --ReAsm` (Section 4.3.2), most of the parental data of the 47 samples are already aligned to GRCh38. Only the parental data of HG002, HG005, and NA21309 were not already aligned, and instead were provided as raw reads.

For these, we used BWA MEM to align the reads to GRCh38. Reads aligning to the IGH locus were then extracted using `samtools view`. Only these reads were provided to the hifiasm trio-binning method, saving time and space compared to if all reads had been provided to hifiasm.

When building the personal reference of the HPRC samples (section 4.3.3), three samples HG01891, HG03453, and NA18906 could not be automatically reconciled, as discussed in Results. We manually reconciled these haplotypes.

In the polishing stage (Section 4.3.5), MaSuRCA used JASPER (30) for read correction. By default, JASPER uses 4 as its threshold for considering a *k*-mer to be “solid,” i.e. unaffected by sequencing error. However, three individuals – HG01952, HG02145, and HG02572 – had coverage histograms that were shifted lower, and so required that we set this parameter to a lower value. In these cases, we set this parameter to 2.

## 5 Acknowledgments

We thank Justin Wagner and Justin Zook of the Genome In A Bottle project for their advice. We thank Kuan-Hao Chao for advice on genome assembly. And we thank Mohsen Zakeri and Sam Kovaka for their help in running the experiments. We are grateful to Corey T. Watson for useful comments.

## 6 Funding

ML and BL were supported by NIGMS grant R35GM139602 and NHGRI grant R01HG011392 to BL.

This work was carried out at the Advanced Research Computing at Hopkins (ARCH) core facility (rockfish.jhu.edu), which is supported by the National Science Foundation (NSF) grant number OAC 1920103.

## 7 Availability of data and materials

The personal assemblies, HiFi raw read, and read alignment of the 47 HPRC samples, and the samples’ parental illumina read, and read alignment are downloaded from the public HPRC S3 bucket https://s3-us-west-2.amazonaws.com/human-pangenomics/index.html?prefix=working/HPRC/ and https://s3-us-west-2.amazonaws.com/human-pangenomics/index.html?prefix=working/HPRC_PLUS/.

The software of IGLoo is available at https://github.com/maojanlin/IGLoo under the MIT license.

## 8 Authors’ contributions

ML and YS designed the method. ML wrote the software and performed the experiments. ML wrote the manuscript. All authors edited and approved the final manuscript.

## 9 Ethics approval

Not applicable.

## 10 Consent for publication

Not applicable.

## 11 Competing interests

The authors declare that they have no competing interests.

## 12 Additional Files

### 12.1 Additional file 1 — Supplementary information

Contains Figures S1–S14.

## Supplementary Material

**Figure S1:**
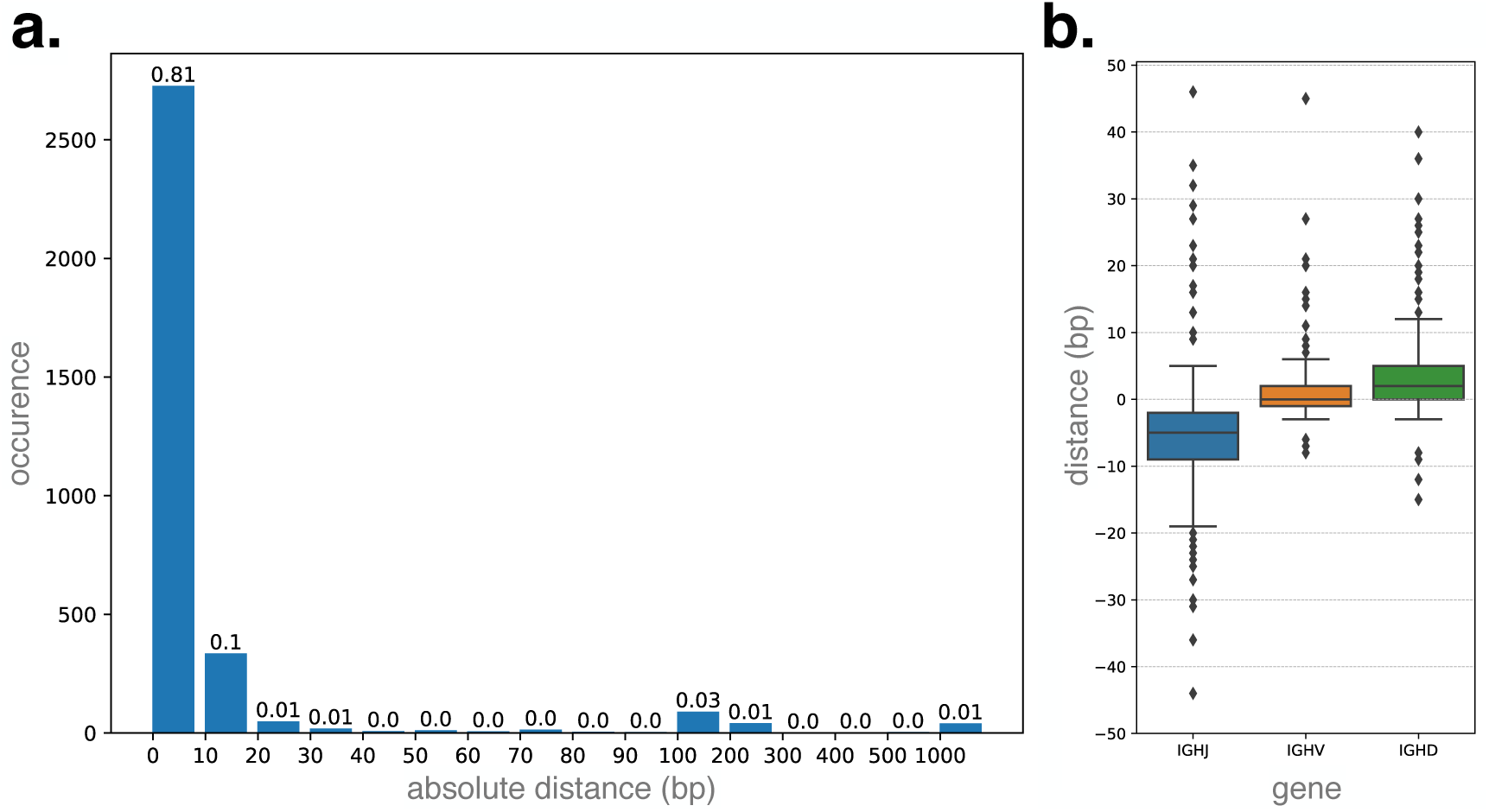
The distribution of the distance of the 3, 357 split site to RSSs. **a.** the histogram of the distribution of the absolute distance (bp) to the closest RSS, the first bin shows the occurrence of distance between 0 to 10, while the last bin shows the occurrence of the distance over 1000. **b.** the distribution of the distance (bp) to RSS stratified by IGHJ, IGHV, and IGHD genes. We only show the cases with distance *<* 50 bp. The distance of J genes are counted on its 5*^′^* end, while the distance of V and D genes are counted from their 3*^′^* end.

**Figure S2:**
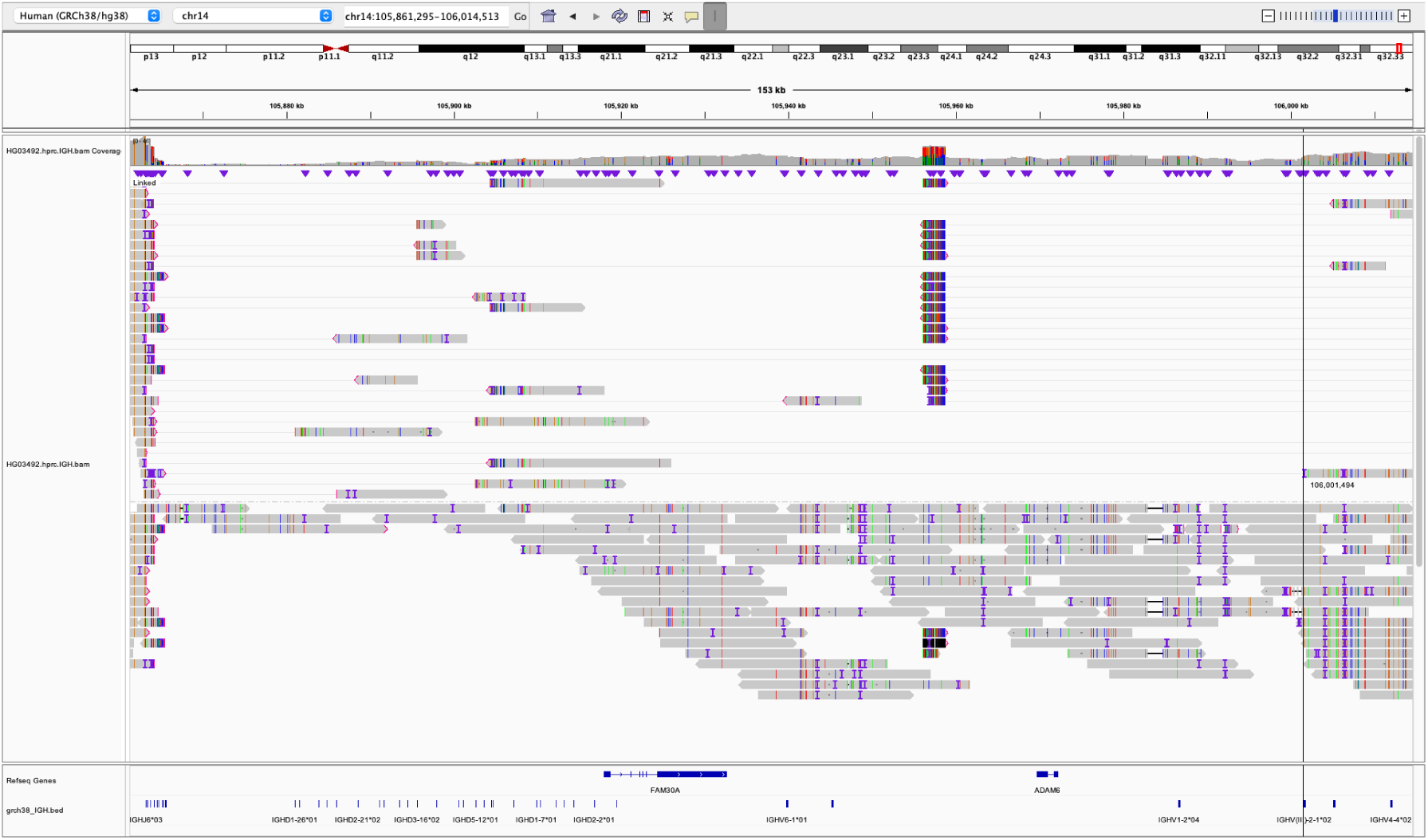
The complete V(D)J recombination event using the pseudogene *IGHV(III)-2-1*.

**Figure S3:**
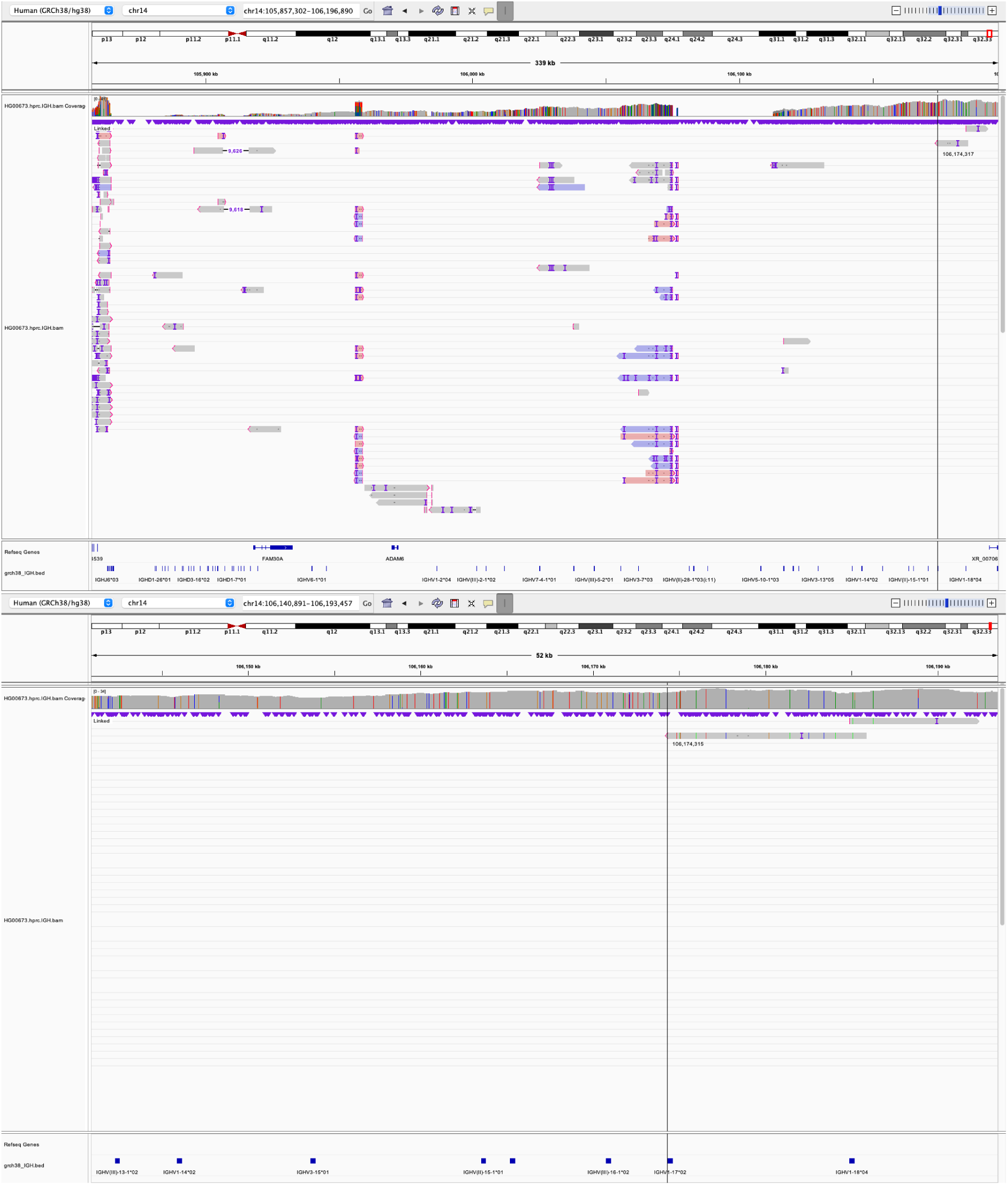
The bird eye view (**top**) and zoom in view (**bottom**) of the complete V(D)J recombination event using the pseudogene *IGHV1-17*.

**Figure S4:**
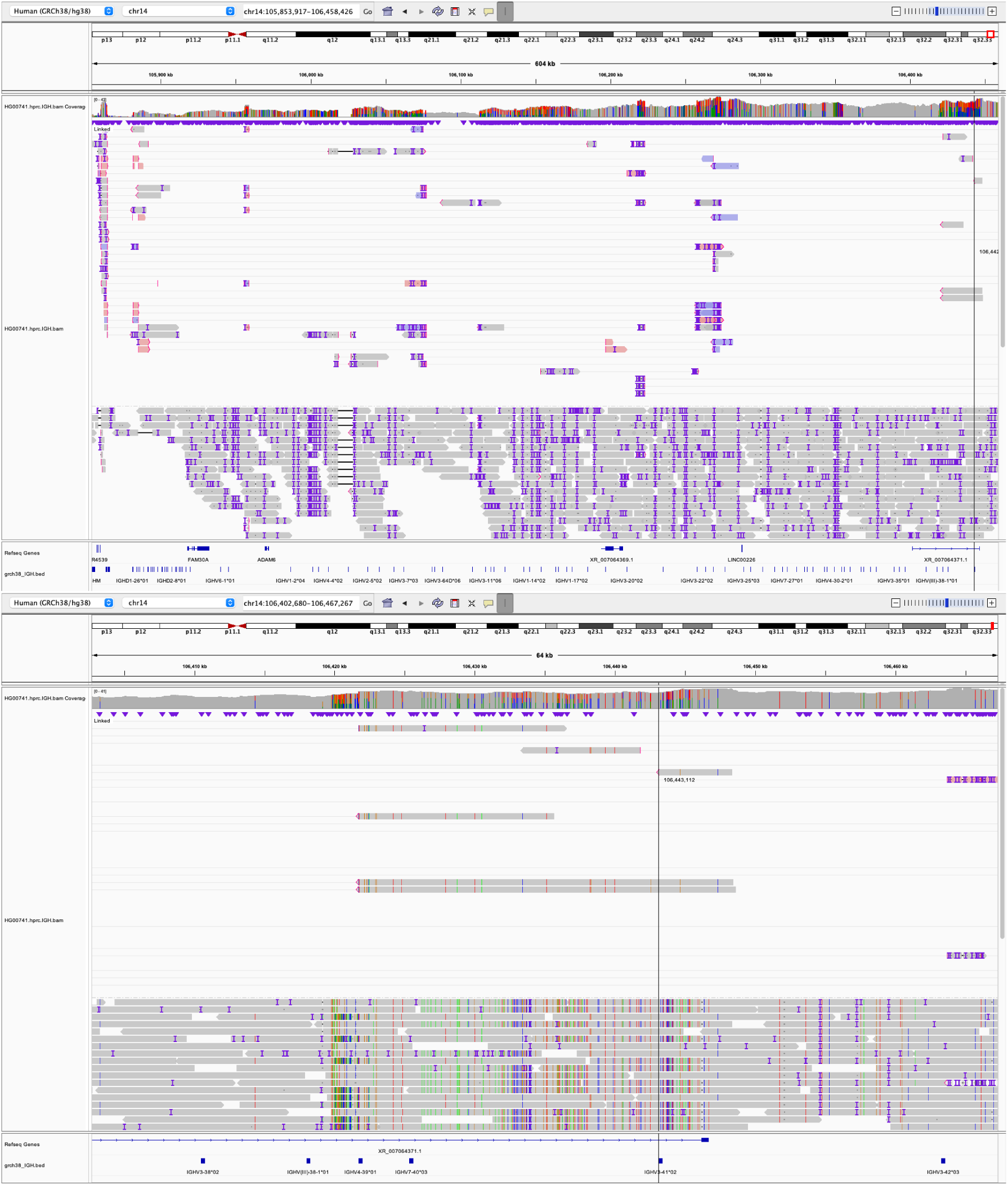
The bird eye view (**top**) and zoom in view (**bottom**) of the complete V(D)J recombination event using the pseudogene *IGHV3-41*.

**Figure S5:**
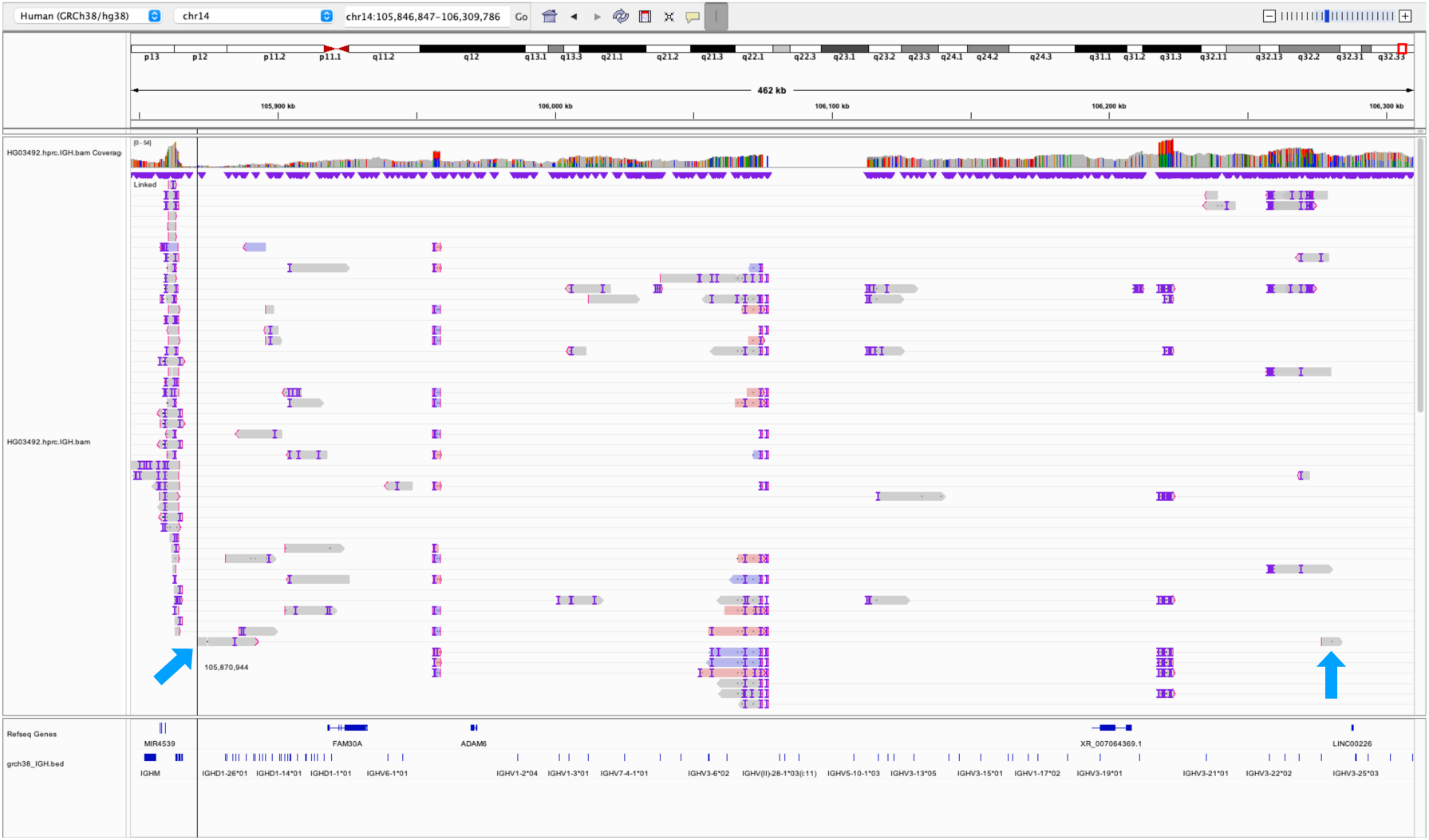
The only V-D only partial recombination event we observed in HPRC from individual HG03492. The blue arrows indicate the reads with a long deletion across the gene *IGHD6-19* and *IGHV1-24*. Note that the 3*^′^* end segments of the read stretch to the position between IGHD and IGHJ loci.

**Figure S6:**
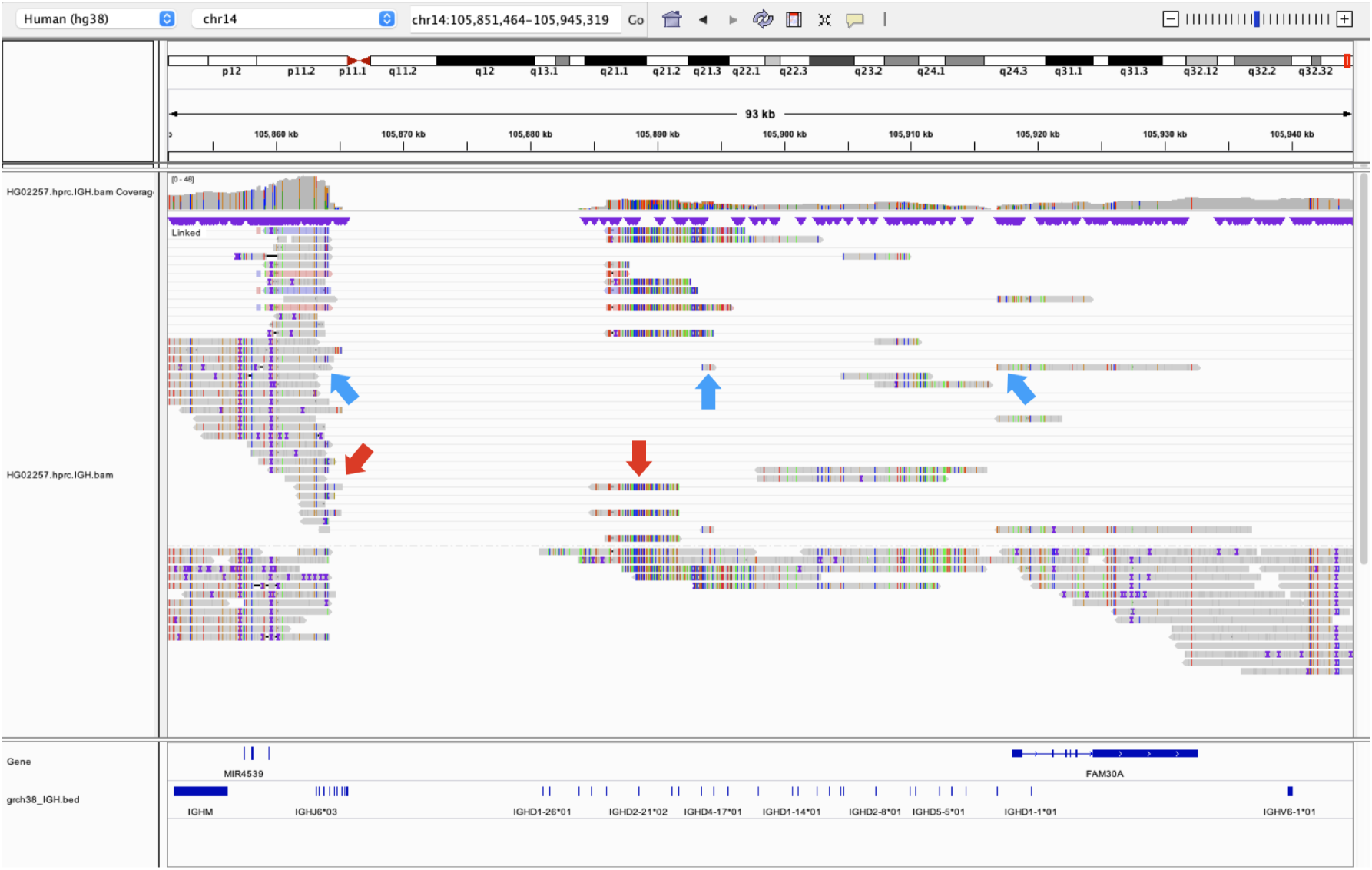
Two non-canonical recombination events of the individual HG02557. The blue arrows indicate the read with two long deletions crossing (*IGHJ4*, *IGHD5-18*) and (*IGHD4-17*, *IGHD2-2*). The red arrows show another read with two long deletions crossing (*IGHJ2*, *IGHD4-23*) and (*IGHD6-19*, *IGHV3-49*). The *IGHV3-49* end is beyond the scope of the screenshot.

**Figure S7:**
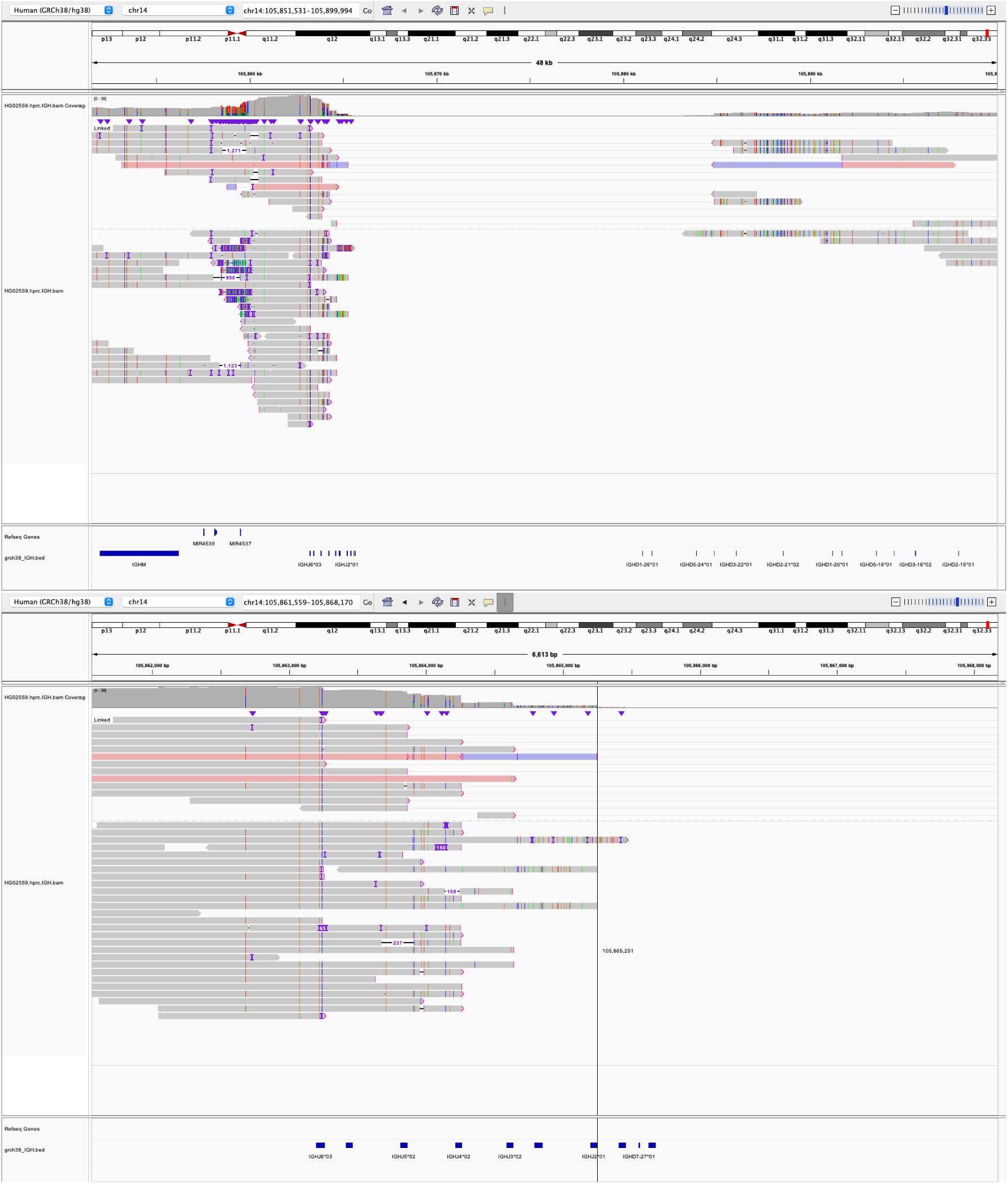
The bird eye view and the zoom in view of the individual HG02559 with an inverted J sequence event. The non-canonical recombination event involved an inversion between *IGHJ2* and *IGHJ4*, the sequence in between the two genes are inverted as the red and blue sequence shown.

**Figure S8:**
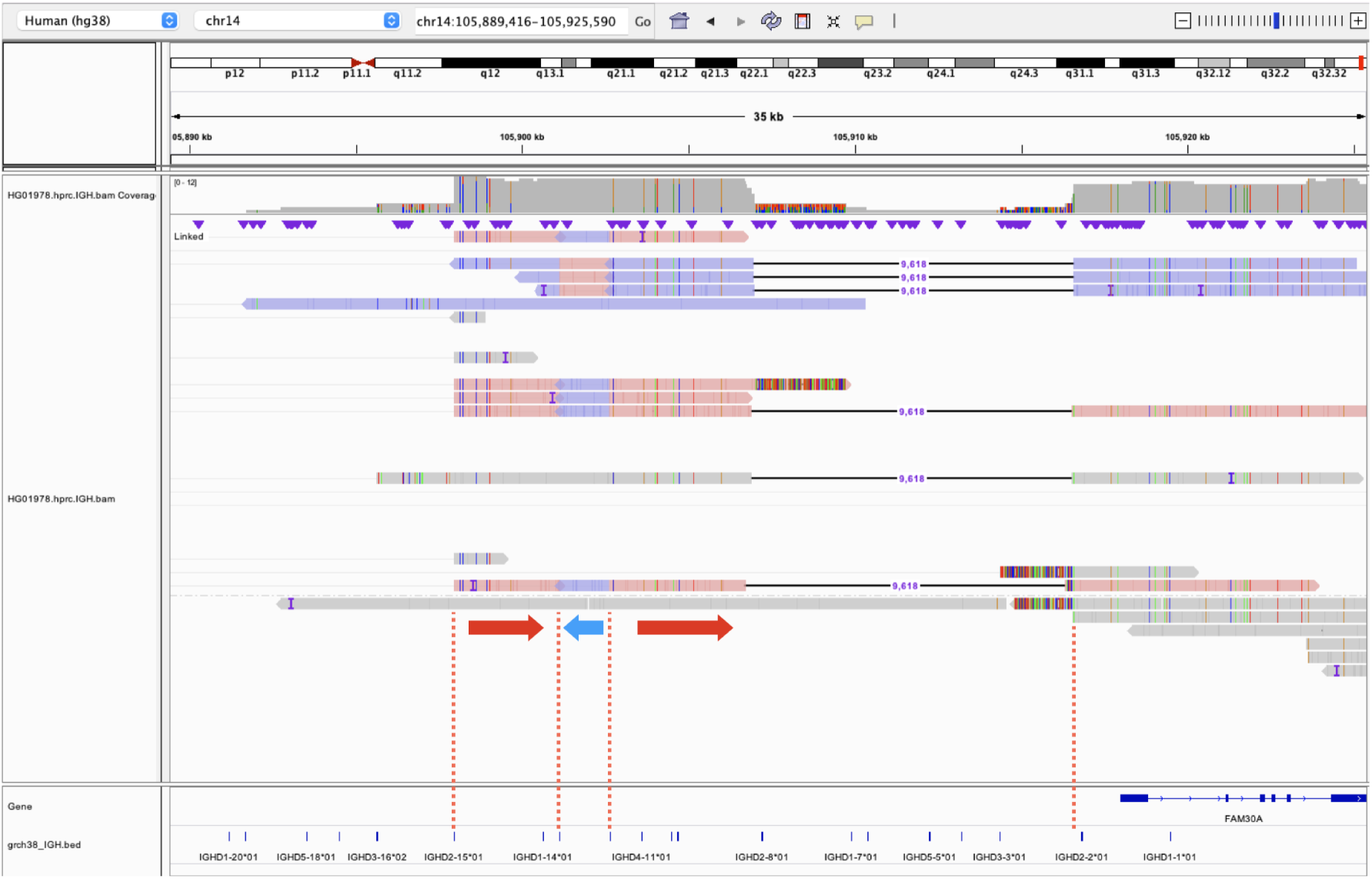
The non-canonical recombination event that involved an inversion between *IGHD6-13* and *IGHD5-12*. The sequence in between the two genes, which indicated by the blue arrow, is inverted. In the raw read, the head of the left side red arrow connects to the tail of the blue arrow, and the blue arrowhead connects to the tail of the right side red arrow. The *IGHD6-13* gene is on the arrowtail of the blue arrow, which is right next to the *IGHD5-12* gene on raw read.

**Figure S9:**
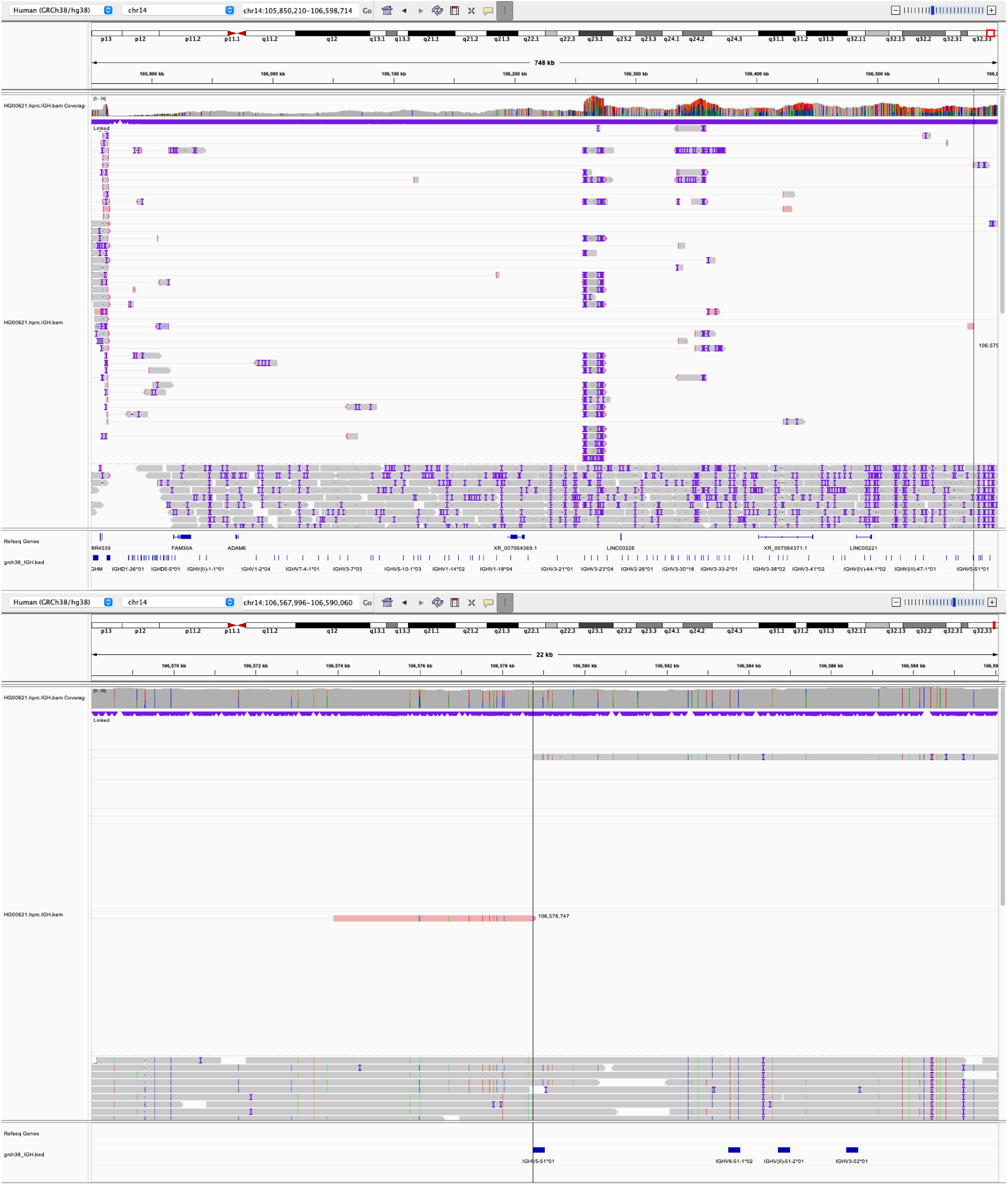
The bird eye view and the zoom in view of the individual HG00621 with an inverted V sequence event. The non-canonical recombination event involved an inversion starting from V gene *IGHV5-51*.

**Figure S10:**
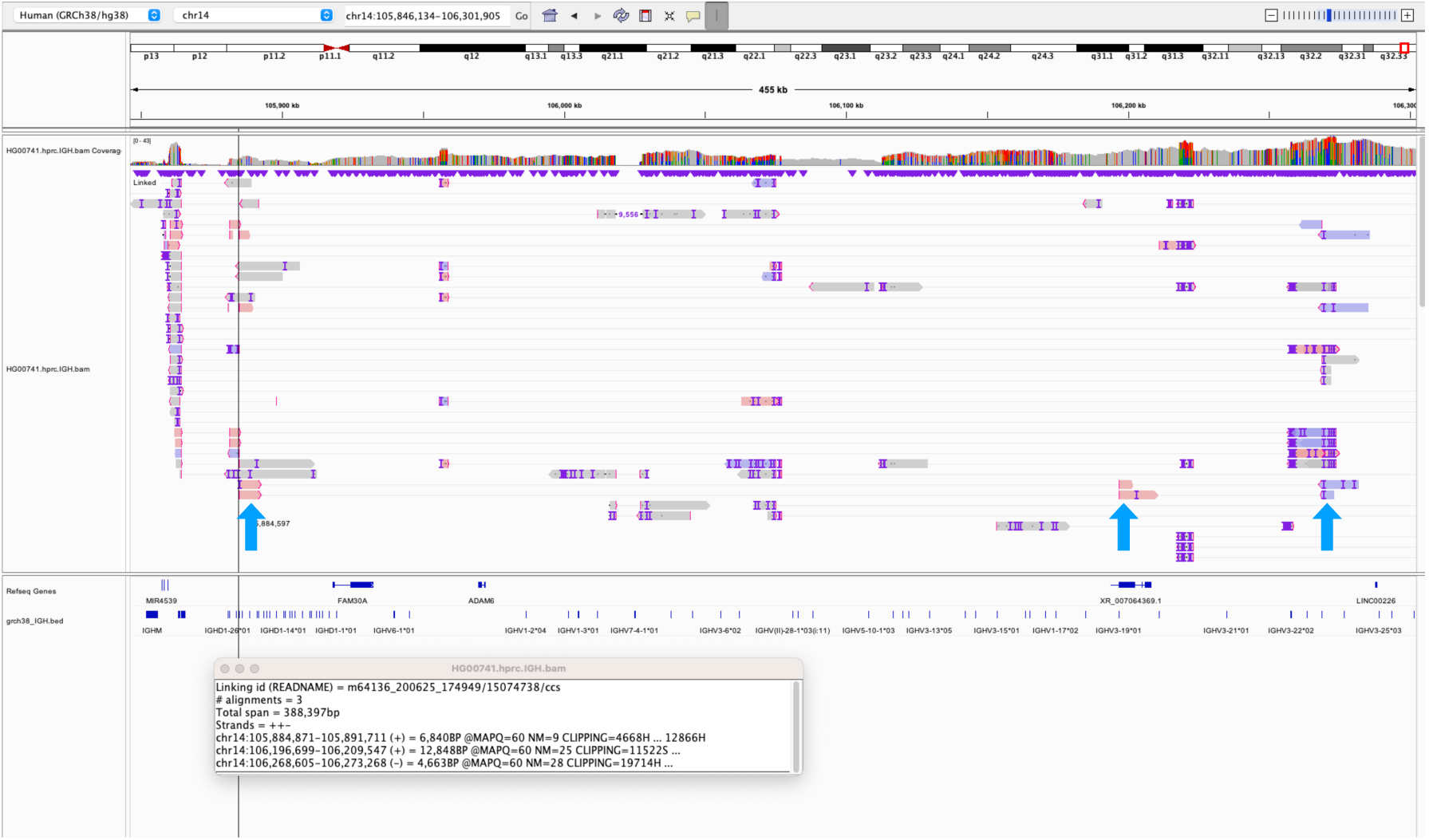
The IGV screenshot of the individual HG00741 with the double V-D recombination event. The segments of the non-canonical recombination event are marked with blue arrows. The event involved a V-D recombination between *IGHD6-19* and *IGHV3-19*, and another V-D recombination event between *IGHD4-23* and *IGHV3-23*.

**Figure S11:**
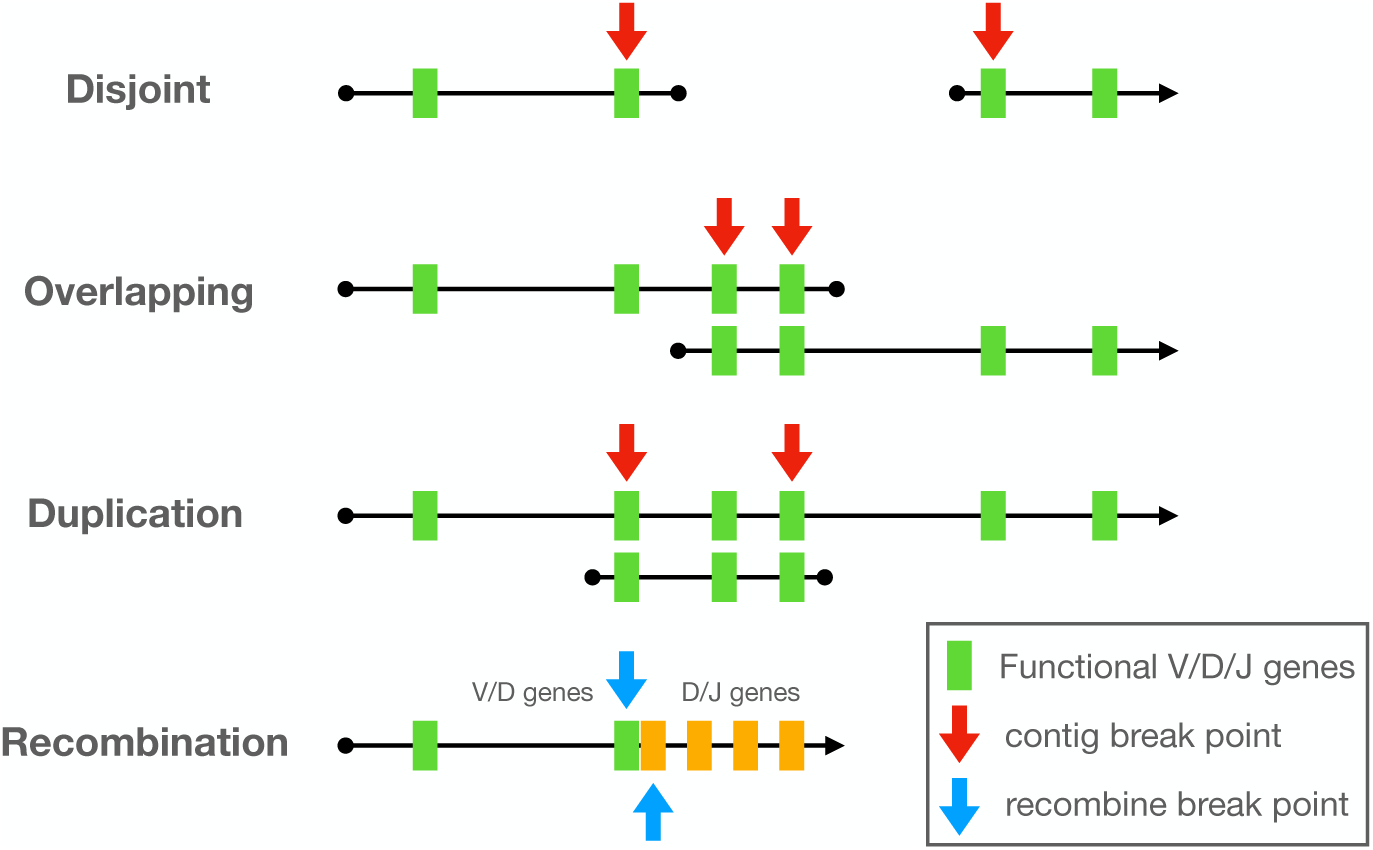
Categories of the breakpoints in IGH locus. **Disjoint**: the contigs end without overlapping each other. **Overlapping**: the assembler reports two separate contigs but with part of the contig overlapping each other. **Duplication**: one contig is contained in the other contig. **Recombination**: the genes changing from V to D/J, or from D to J with non-germline transition.

**Figure S12:**
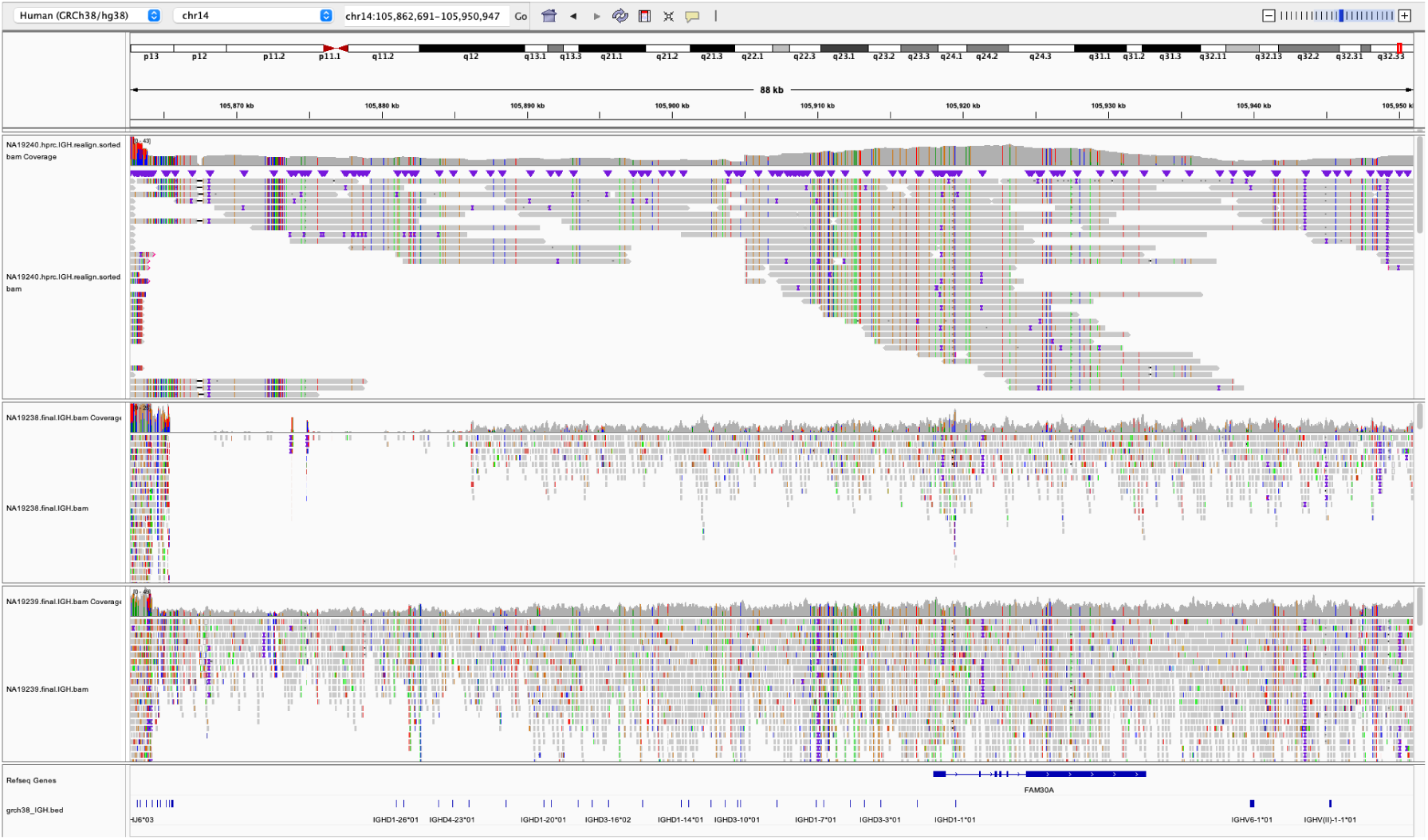
The IGV screenshot of the read mapping on the IGHD loci of the sample NA19240 and the sample’s maternal (NA19238) and paternal (NA19239) data.

**Figure S13:**
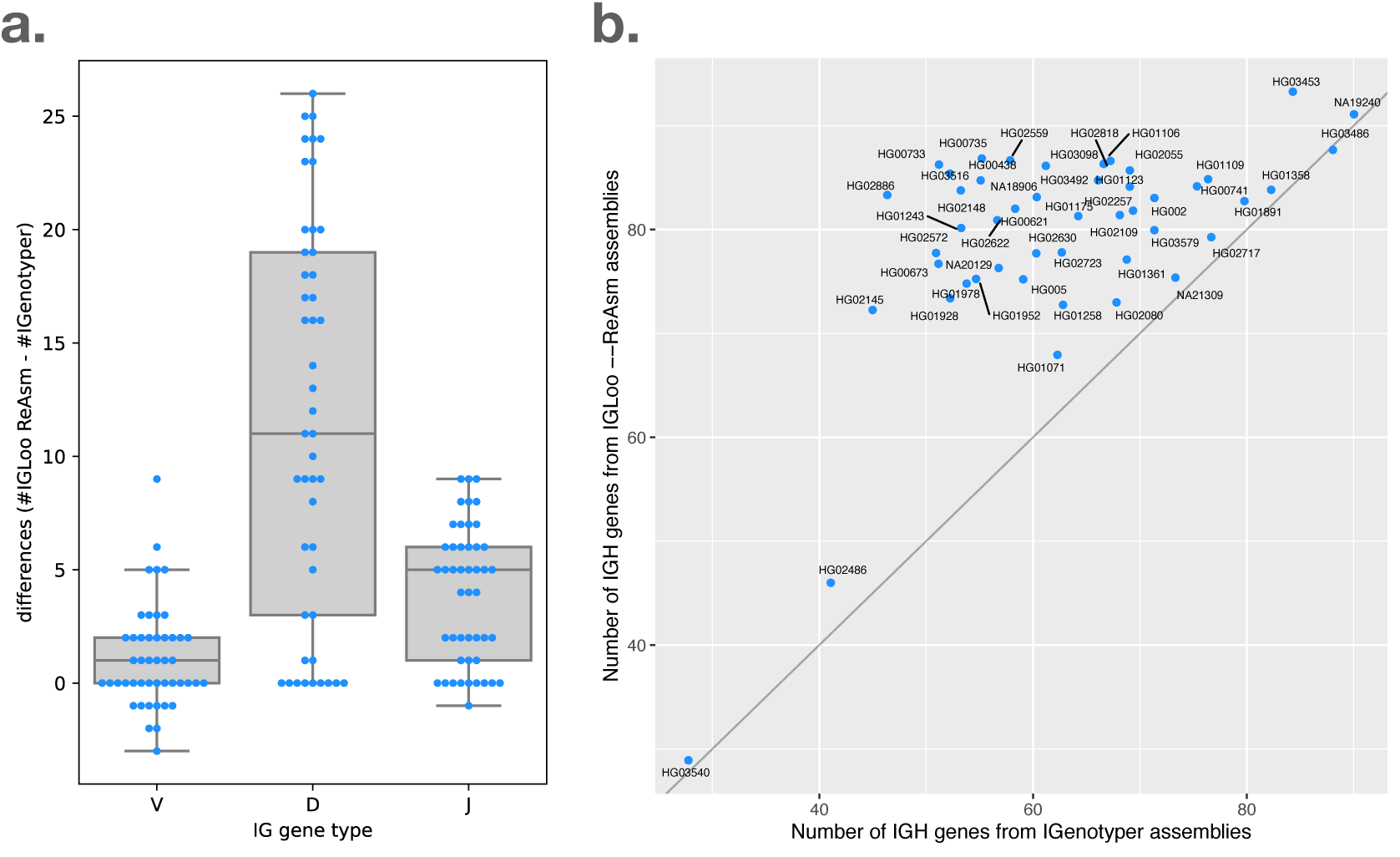
The IGH gene number comparison between IGenotyper (14) and `IGLoo --ReAsm` in the 47 HPRC samples. **a.** the V, D, and J gene differences between two methods in IGH locus. The positive value means `IGLoo --ReAsm` assemblies cover more IGH genes while the negative value means IGenotyper assemblies cover more genes. **b.** the total number of IGH V, D, and J genes comparison of the two methods.

**Figure S14:**
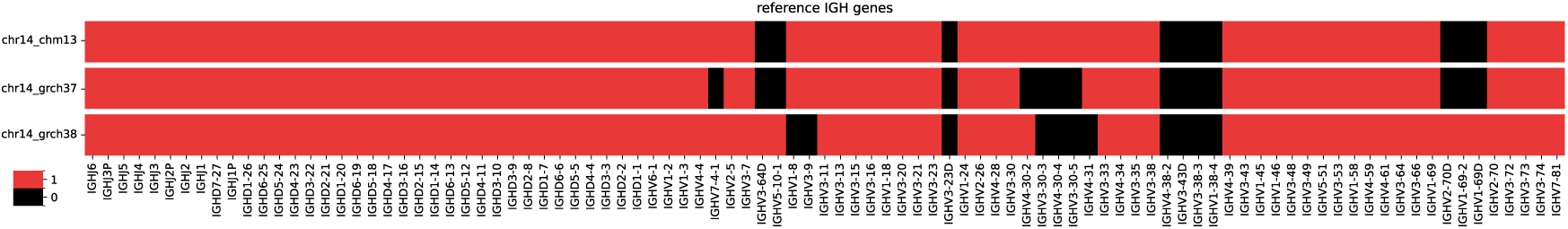
the gene number in its relative position on the chromosome 14 of reference genomes T2T-CHM13, GRCh37, and GRCh38. All D and J genes are listed, but only functional or ORF V genes are listed for simplicity

## Notes

### Competing Interest Statement

The authors have declared no competing interest.

## References

[1] Susumu Tonegawa. Somatic generation of antibody diversity. Nature, 302(5909):575– 581, 1983.

[2] Martin Fuxa, Jane Skok, Abdallah Souabni, Giorgia Salvagiotto, Esther Roldan, and Meinrad Busslinger. Pax5 induces v-to-dj rearrangements and locus contraction of the immunoglobulin heavy-chain gene. Genes & development, 18(4):411–422, 2004.

[3] Oscar L Rodriguez, Yana Safonova, Catherine A Silver, Kaitlyn Shields, William S Gibson, Justin T Kos, David Tieri, Hanzhong Ke, Katherine JL Jackson, Scott D Boyd, et al. Genetic variation in the immunoglobulin heavy chain locus shapes the human antibody repertoire. Nature communications, 14(1):4419, 2023.

[4] Yuval Avnir, Corey T Watson, Jacob Glanville, Eric C Peterson, Aimee S Tallarico, Andrew S Bennett, Kun Qin, Ying Fu, Chiung-Yu Huang, John H Beigel, et al. Ighv1-69 polymorphism modulates anti-influenza antibody repertoires, correlates with ighv utilization shifts and varies by ethnicity. Scientific reports, 6(1):20842, 2016.

[5] Jeong Hyun Lee, Laura Toy, Justin T Kos, Yana Safonova, William R Schief, Colin Havenar-Daughton, Corey T Watson, and Shane Crotty. Vaccine genetics of ighv1-2 vrc01-class broadly neutralizing antibody precursor näıve human b cells. NPJ vaccines, 6(1):113, 2021.

[6] Tabish Hussain and Rita Mulherkar. Lymphoblastoid cell lines: a continuous in vitro source of cells to study carcinogen sensitivity and dna repair. International journal of molecular and cellular medicine, 1(2):75, 2012.

[7] Dorothee Nickles, Lohith Madireddy, Shan Yang, Pouya Khankhanian, Steve Lincoln, Stephen L Hauser, Jorge R Oksenberg, and Sergio E Baranzini. In depth comparison of an individual’s dna and its lymphoblastoid cell line using whole genome sequencing. BMC genomics, 13:1–11, 2012.

[8] Nayanah Siva. 1000 genomes project. Nature biotechnology, 26(3):256–257, 2008.

[9] Richard A Gibbs, John W Belmont, Paul Hardenbol, Thomas D Willis, Fuli L Yu, HM Yang, Lan-Yang Ch’ang, Wei Huang, Bin Liu, Yan Shen, et al. The international hapmap project. 2003.

[10] Justin M Zook, David Catoe, Jennifer McDaniel, Lindsay Vang, Noah Spies, Arend Sidow, Ziming Weng, Yuling Liu, Christopher E Mason, Noah Alexander, et al. Extensive sequencing of seven human genomes to characterize benchmark reference materials. Scientific data, 3(1):1–26, 2016.

[11] Ting Wang, Lucinda Antonacci-Fulton, Kerstin Howe, Heather A Lawson, Julian K Lucas, Adam M Phillippy, Alice B Popejoy, Mobin Asri, Caryn Carson, Mark JP Chaisson, et al. The human pangenome project: a global resource to map genomic diversity. Nature, 604(7906):437–446, 2022.

[12] Corey T Watson, Karyn M Steinberg, John Huddleston, Rene L Warren, Maika Malig, Jacqueline Schein, A Jeremy Willsey, Jeffrey B Joy, Jamie K Scott, Tina A Graves, et al. Complete haplotype sequence of the human immunoglobulin heavy-chain variable, diversity, and joining genes and characterization of allelic and copy-number variation. The American Journal of Human Genetics, 92(4):530–546, 2013.

[13] Oscar L Rodriguez, Andrew J Sharp, and Corey T Watson. Limitations of lymphoblastoid cell lines for establishing genetic reference datasets in the immunoglobulin loci. Plos one, 16(12):e0261374, 2021.

[14] Oscar L Rodriguez, William S Gibson, Wayne A Marasco, Robert Sebra, Melissa L Smith, and Corey T Watson. A novel framework for characterizing genomic haplotype diversity in the human immunoglobulin heavy chain locus. Frontiers in immunology, 11:571270, 2020.

[15] William S Gibson, Oscar L Rodriguez, Kaitlyn Shields, Catherine A Silver, Abdullah Dorgham, Matthew Emery, Gintaras Deikus, Robert Sebra, Evan E Eichler, Ali Bashir, et al. Characterization of the immunoglobulin lambda chain locus from diverse populations reveals extensive genetic variation. Genes & Immunity, 24(1):21–31, 2023.

[16] Haoyu Cheng, Gregory T Concepcion, Xiaowen Feng, Haowen Zhang, and Heng Li. Haplotype-resolved de novo assembly using phased assembly graphs with hifiasm. Nature methods, 18(2):170–175, 2021.

[17] Aleksey V Zimin, Guillaume Marçais, Daniela Puiu, Michael Roberts, Steven L Salzberg, and James A Yorke. The masurca genome assembler. Bioinformatics, 29(21):2669–2677, 2013.

[18] Yana Safonova and Pavel A Pevzner. De novo inference of diversity genes and analysis of non-canonical v (dd) j recombination in immunoglobulins. Frontiers in immunology, 10:448987, 2019.

[19] Joshua Tan, Kathrin Pieper, Luca Piccoli, Abdirahman Abdi, Mathilde Foglierini, Roger Geiger, Claire Maria Tully, David Jarrossay, Francis Maina Ndungu, Juliana Wambua, et al. A lair1 insertion generates broadly reactive antibodies against malaria variant antigens. Nature, 529(7584):105–109, 2016.

[20] Heng Li. Minimap2: pairwise alignment for nucleotide sequences. Bioinformatics, 34(18):3094–3100, 2018.

[21] D. M. Church, V. A. Schneider, T. Graves, K. Auger, F. Cunningham, N. Bouk, H. C. Chen, R. Agarwala, W. M. McLaren, G. R. Ritchie, D. Albracht, M. Kremitzki, S. Rock, H. Kotkiewicz, C. Kremitzki, A. Wollam, L. Trani, L. Fulton, R. Fulton, L. Matthews, S. Whitehead, W. Chow, J. Torrance, M. Dunn, G. Harden, G. Threadgold, J. Wood, J. Collins, P. Heath, G. Griffiths, S. Pelan, D. Grafham, E. E. Eichler, G. Weinstock, E. R. Mardis, R. K. Wilson, K. Howe, P. Flicek, and T. Hubbard. Modernizing reference genome assemblies. PLoS Biol., 9(7):e1001091, Jul 2011.

[22] D. M. Church, V. A. Schneider, K. M. Steinberg, M. C. Schatz, A. R. Quinlan, C. S. Chin, P. A. Kitts, B. Aken, G. T. Marth, M. M. Hoffman, J. Herrero, M. L. Mendoza, R. Durbin, and P. Flicek. Extending reference assembly models. Genome Biol., 16:13, 1 2015.

[23] S. Nurk, S. Koren, A. Rhie, M. Rautiainen, A. V. Bzikadze, A. Mikheenko, M. R. Vollger, N. Altemose, L. Uralsky, A. Gershman, S. Aganezov, S. J. Hoyt, M. Diekhans, G. A. Logsdon, M. Alonge, S. E. Antonarakis, M. Borchers, G. G. Bouffard, S. Y. Brooks, G. V. Caldas, N. C. Chen, H. Cheng, C. S. Chin, W. Chow, L. G. de Lima, P. C. Dishuck, R. Durbin, T. Dvorkina, I. T. Fiddes, G. Formenti, R. S. Fulton, A. Fungtammasan, E. Garrison, P. G. S. Grady, T. A. Graves-Lindsay, I. M. Hall, N. F. Hansen, G. A. Hartley, M. Haukness, K. Howe, M. W. Hunkapiller, C. Jain, M. Jain, E. D. Jarvis, P. Kerpedjiev, M. Kirsche, M. Kolmogorov, J. Korlach, M. Kremitzki, H. Li, V. V. Maduro, T. Marschall, A. M. McCartney, J. McDaniel, D. E. Miller, J. C. Mullikin, E. W. Myers, N. D. Olson, B. Paten, P. Peluso, P. A. Pevzner, D. Porubsky, T. Potapova, E. I. Rogaev, J. A. Rosenfeld, S. L. Salzberg, V. A. Schneider, F. J. Sedlazeck, K. Shafin, C. J. Shew, A. Shumate, Y. Sims, A. F. A. Smit, D. C. Soto, I. ć, J. M. Storer, A. Streets, B. A. Sullivan, F. Thibaud-Nissen, J. Torrance, J. Wagner, B. P. Walenz, A. Wenger, J. M. D. Wood, C. Xiao, S. M. Yan, A. C. Young, S. Zarate, U. Surti, R. C. McCoy, M. Y. Dennis, I. A. Alexandrov, J. L. Gerton, R. J. O’Neill, W. Timp, J. M. Zook, M. C. Schatz, E. E. Eichler, K. H. Miga, and A. M. Phillippy. The complete sequence of a human genome. Science, 376(6588):44–53, Apr 2022.

[24] Marie-Paule Lefranc, Veronique Giudicelli, Chantal Ginestoux, Joumana Jabado-Michaloud, Geraldine Folch, Fatena Bellahcene, Yan Wu, Elodie Gemrot, Xavier Brochet, Jerôme Lane, et al. Imgt®, the international immunogenetics information system®. Nucleic acids research, 37(suppl 1):D1006–D1012, 2009.

[25] Mao-Jan Lin, Yu-Chun Lin, Nae-Chyun Chen, Allen Chilun Luo, Sheng-Kai Lai, Chia-Lang Hsu, Jacob Shujui Hsu, Chien-Yu Chen, Wei-Shiung Yang, and Pei-Lung Chen. Profiling genes encoding the adaptive immune receptor repertoire with gairr suite. Frontiers in Immunology, 13:922513, 2022.

[26] Alexander P Sweeten, Michael C Schatz, and Adam M Phillippy. Moddotplot-rapid and interactive visualization of complex repeats. bioRxiv, pages 2024–04, 2024.

[27] Heng Li. Aligning sequence reads, clone sequences and assembly contigs with bwamem. arXiv preprint arXiv:1303.3997, 2013.

[28] S. Koren, A. Rhie, B. P. Walenz, A. T. Dilthey, D. M. Bickhart, S. B. Kingan, S. Hiendleder, J. L. Williams, T. P. L. Smith, and A. M. Phillippy. De novo assembly of haplotype-resolved genomes with trio binning. Nat Biotechnol, Oct 2018.

[29] Ole Tange. Gnu parallel 20220522 (’nato’), May 2022. GNU Parallel is a general parallelizer to run multiple serial command line programs in parallel without changing them.

[30] Alina Guo, Steven L Salzberg, and Aleksey V Zimin. Jasper: A fast genome polishing tool that improves accuracy of genome assemblies. PLoS computational biology, 19(3):e1011032, 2023.

